# Comparison between EEG and MEG of static and dynamic resting-state networks

**DOI:** 10.1101/2024.04.29.591774

**Authors:** SungJun Cho, Mats van Es, Mark Woolrich, Chetan Gohil

**Affiliations:** Oxford Centre for Human Brain Activity, Wellcome Centre for Integrative Neuroimaging, Department of Psychiatry, University of Oxford, United Kingdom, OX3 7JK

## Abstract

The characterisation of resting state networks (RSNs) using neuroimaging techniques has significantly contributed to our understanding of the organisation of brain activity. Prior work has demonstrated the electrophysiological basis of RSNs and their dynamic nature, revealing transient activations of brain networks with millisecond timescales. While previous research has confirmed the comparability of RSNs identified by electroencephalography (EEG) to those identified by magnetoencephalography (MEG) and functional magnetic resonance imaging (fMRI), most studies have utilised static analysis techniques, ignoring the dynamic nature of brain activity. Often, these studies use high-density EEG systems, which limit their applicability in clinical settings. Addressing these gaps, our research studies RSNs using medium-density EEG systems (61 sensors), comparing both static and dynamic brain network features to those obtained from a high-density MEG system (306 sensors). We assess the qualitative and quantitative comparability of EEG-derived RSNs to those from MEG, including their ability to capture age-related effects, and explore the reproducibility of dynamic RSNs within and across the modalities. Our findings suggest that both MEG and EEG offer comparable static and dynamic network descriptions, albeit with MEG offering some increased sensitivity and reproducibility. Such RSNs and their comparability across the two modalities remained consistent qualitatively but not quantitatively when the data were reconstructed without subject-specific structural MRI images.

## 1 Introduction

Resting-state networks (RSNs) were first observed in functional magnetic resonance imaging (fMRI) and positron emission tomography (PET) in the late 20th century and later independently identified in magnetoencephalography (MEG) and electroencephalography (EEG) [1, 2]. A study by Brookes et al. (2013) provided robust evidence for the electrophysiological basis of RSNs, employing temporal independent component analysis (ICA) on MEG amplitude envelopes to demonstrate that the spatial patterns of brain networks in MEG resemble those of the RSNs previously identified by fMRI [3]. While early descriptions of RSNs were essentially static (i.e., time-averaged), further research has revealed that they can be well described as activating in a transient manner, with different networks activating at distinct time points [4, 5]. The contribution of electrophysiology to our understanding of RSNs is underscored by their functional significance. MEG and EEG studies have allowed RSNs to be linked to sensorimotor functions at fast sub-second timescales, as well as to a range of cognitive tasks concerning working memory and language comprehension [6–8]. Research on neuropsychiatric disorders has also elucidated the functional relevance of RSNs across various pathological conditions, including Alzheimer’s disease, major depressive disorder, and schizophrenia, highlighting their utility in understanding brain dysfunction [9–11]. From a technical standpoint, electrophysiological analysis of RSNs offers unparalleled insight into the spectral and temporal domains. M/EEG-derived functional networks allow for the investigation of their spectral contents; empirical evidence suggests that neural activities can be segmented into distinct states of global synchronous fluctuations, each characterised by specific oscillatory frequencies [12, 13]. Furthermore, the superior temporal resolution of electrophysiological approaches facilitates a detailed examination of the transient dynamics of resting-state and task-based networks. For example, previous research indicates that M/EEG-derived RSNs activate on faster timescales (*∼*100-200 ms) than those discerned through fMRI (*∼*10 s) [4, 5].

To date, studies examining electrophysiological RSNs, particularly those based on the amplitude time-courses of brain regions, have affirmed a reasonable comparability of EEG RSNs to their MEG or fMRI counterparts [14–17]. However, these studies relied on static analyses (by which we mean that they do not provide a time-varying description of interactions between brain areas), such as spatial ICA or seed-based correlation. More recently, Coquelet et al. (2020) evaluated the correspondence of RSNs between 306-channel MEG and 256-channel EEG data [18]. This study revealed that while static functional connectivity (FC) patterns are generally consistent across modalities in the resting state, dynamic FC shows less agreement. Their dynamic FC methods, however, used sliding windows and clustering approaches, which are heavily dependent on user-defined hyperparameters and compromise temporal resolution. Finally, studies to date have typically employed high-density EEG systems (*>* 100 channels) [15, 17, 18], facilitating the computation of RSNs in brain space, where spatial leakage can be more appropriately controlled and the networks are more interpretable [19]. Yet, this overlooks the wider availability and clinical utility of lower-density EEG systems, for which the ability to reliably estimate RSNs in brain space would be a valuable asset.

Acknowledging these limitations, this paper aims to establish the efficacy of EEG in providing static and dynamic functional network perspectives akin to those obtained from MEG and fMRI within brain space, with a state-of-the-art dynamic modelling technique—Time-Delay Embedded Hidden Markov Model (TDE-HMM) [20] and without the need for high density data. Drawing on existing literature, we hypothesised that both EEG and MEG will provide comparable static and dynamic RSN descriptions, with dynamic features anticipated to exhibit less consistency between modalities. Using MEG RSNs as a benchmark for assessing fast oscillatory networks, we compared static and dynamic network descriptions against those derived from EEG, with the aim of establishing a basis set of EEG RSNs that can be identified with medium-density EEG data (*∼*61 channels). In addition, we also examined whether M/EEG RSNs, derived from data reconstructed without subject-specific structural MRI (sMRI) images, can yield network descriptions similar to those obtained using subject sMRI images without loss of quality.

This endeavour seeks to demonstrate that MEG and EEG provide similar and reliable network descriptions, allowing for research that can leverage the accessibility and cost-effectiveness of EEG in the exploration of brain oscillatory networks in both health and disease. Additionally, this work provides a publicly available set of M/EEG RSNs, complete with scripts for generating precise network inferences, thereby contributing to our comprehensive understanding of the complementary potential of MEG and EEG in brain network research.

## 2 Methods

### 2.1 Datasets

In this study, we used the openly available resting-state EEG and MEG datasets called the Leipzig Study for Mind-Body-Emotion Interactions (LEMON) dataset and the Cambridge Centre for Ageing and Neuroscience (Cam-CAN) dataset, respectively. Both datasets were obtained from healthy adults, and recordings were measured while participants were seated. The two datasets were matched to have an identical age distribution and number of subjects. To specify, each dataset was divided into subsets of a 5-year age interval, and for imbalances between each subset of two datasets, subjects were randomly subsampled from the larger subset to ensure parity. The age-matched datasets comprised a total of 96 subjects, consisting of 60 young (aged between 20-35 years) and 36 old (aged between 55-80 years) participants^1^ (Fig. A1). In the EEG data, the young group comprised 44 females and 16 males, while the old group consisted of 23 females and 13 males. In the MEG data, the young group comprised 25 females and 35 males, while the old group consisted of 19 females and 17 males. The protocols and demographics related to the data collection are outlined in [21] for LEMON and [22] for Cam-CAN in detail and are partly repeated here for clarity.

#### 2.1.1 EEG Data

The EEG recordings were obtained using a BrainAMP MR plus amplifier with 62-channel ActiCAP electrodes (Brain Products, Gilchling, Germany). These channels consisted of 61 EEG electrodes and 1 vertical electrooculogram (VEOG) channel. A surrogate horizontal electrooculogram (HEOG) channel was additionally added by taking the difference between channels F7 and F8. The channel montage adapted the 10-10 layout, which was referenced and grounded at FCz and the sternum, respectively. The acquisition time of the recordings was 16 min, with 1 min blocks alternating between eyes-closed and eyes-open conditions. Signals were collected with a sampling rate of 2500 Hz and bandpass filtered between 0.015 Hz and 1 kHz. For coregistration purposes, the T1-weighted sMRI was collected over 8 min 22 s, using Magnetization-Prepared 2 RApid Acquisition Gradient Echos (MP2RAGE) sequences with a 3T MAGNETOM Verio scanner (Siemens Healthcare, Erlangen, Germany) equipped with a 32-channel head coil.

#### 2.1.2 MEG Data

The MEG recordings were obtained using a 306-channel Vector View system (Elekta Neuromag, Helsinki, Finland), consisting of 102 magnetometers and 204 orthogonal planar gradiometers. The acquisition time of the recordings was 8 min 40 s, with the first 20 s discarded. Only eyes-closed resting-state data were recorded at a sampling rate of 1 kHz, bandpass filtered between 0.03 Hz and 330 Hz. The eye- and pulse-related artefacts were monitored using two pairs of biopolar EOG electrodes and one pair of bipolar electrocardiogram (ECG) electrodes, respectively. For coregistration, the T1-weighted sMRI was recorded for 4 min 32 s, using a Magnetization-Prepared RApid Gradient Echos (MPRAGE) sequence with a 3T TIM Trio scanner (Siemens Healthcare, Erlangen, Germany) equipped with a 32-channel head coil.

### 2.2 Data Preprocessing and Source Reconstruction

The preprocessing and source reconstruction processes largely followed the pipeline outlined in [23]. Detailed steps of the pipeline applied to the M/EEG datasets were slightly different due to the unique data characteristics intrinsic to each dataset. However, they were aligned as closely as possible to mitigate any unknown variance that could arise from discrepancies in these steps. Before applying the pipeline, the MEG data were Maxfiltered using the temporal signal-space separation (tSSS) method to separate recordings of neural activity within the brain from any external noise sources [6]. The entire process of preprocessing and source reconstruction was conducted using the OHBA Software Library (OSL) package [24].

#### 2.2.1 Preprocessing

The LEMON and Cam-CAN data were first cropped to exclude the first 30 seconds of the recordings to account for the subject acclimation and stabilisation of measurement devices. These data were band-pass filtered between 0.5 and 125 Hz using a fifth-order IIR Butterworth filter and subsequently notch filtered at 50 and 100 Hz to remove mains artefact (and additionally at 88 Hz for Cam-CAN to remove spike artefacts). The data were then downsampled to 250 Hz. The bad segments and channels with significantly high variance were removed using an automated generalised-extreme studentised deviate (G-ESD) algorithm [25]. For Cam-CAN, bad segment and channel detection procedures were applied separately to different sensor types (i.e., magnetometers and gradiometers). The data were further denoised by applying a FastICA decomposition [26] to the M/EEG data, decomposing signals into 64 and 54 components, respectively. Components associated with ocular artefacts (i.e., blinks and saccades) were removed, and ECG-related artefacts were additionally excluded for Cam-CAN. Prior to the ICA decomposition, bad segments from EOG recordings were identified to prevent the inadvertent removal of pertinent M/EEG signals due to concurrent noise artefacts in both M/EEG and EOG. Finally, to keep the data dimension consistent across the subjects, any bad channels detected earlier were interpolated from ICA-cleaned data using spherical spline interpolation [27].

#### 2.2.2 Source Reconstruction

The M/EEG data were coregistered to sMRI image and digitised head points using the affine transformation algorithm in OSL. For LEMON, a boundary element model (BEM) with triple layers of scalp, skull, and cortex surfaces was employed as a head model; for Cam-CAN, a BEM with a single layer of a scalp surface was employed. The surfaces of a scalp, inner skull, and brain were extracted from sMRI data using FSL’s BET tool [28, 29]. The nose was not included during the coregistration step, since the sMRI images were defaced.

After coregistration, the preprocessed sensor data were band-pass filtered between 1-45 Hz and reconstructed onto an 8 mm isotropic grid using a linearly constrained minimum variance (LCMV) volumetric beamformer [30]. The rank of a data covariance matrix used to compute the beamformer weights was set to 50^2^, which regularises the covariance estimation and is comfortably below the rank of both the MEG (following Maxfiltering) and EEG data. Voxels were then parcellated into 38 anatomically defined regions. Source reconstructed signals were obtained by applying principal component analysis (PCA) to these voxel-wise data, wherein a time series of each parcel would be the first principal component that explains most variance from all voxels within a parcel. The symmetric multivariate leakage reduction technique [19] was applied to minimise spurious correlations between all parcels and reduce source/spatial leakage including so-called ‘inherited’ or ‘ghost interactions’ [31]. Lastly, the combination of the LCMV volumetric beamformer and the PCA used in the parcel time series calculations means that the sign of the estimated parcel time series varies arbitrarily across subjects. The sign of each channel was therefore matched across subjects using a random flip algorithm described in [20]. The source reconstruction procedure was repeated using the standard Montreal Neurological Institute (MNI) T1 structural brain image instead of subject-specific sMRI images. We employed this approach to explore whether similar M/EEG RSN features could be discerned using a standard brain structural image (i.e., without subject sMRI files) and if the comparability between MEG and EEG remains under these conditions. Unless indicated otherwise, all results presented herein are based on the data reconstructed *with* the subject-specific sMRI images.

#### 2.2.3 Data Organisation

To make the LEMON and Cam-CAN dataset more comparable, only eyes-closed segments were extracted from the source reconstructed EEG LEMON data. After extracting the eyes-closed segments, the mean and standard deviation of the data lengths were 476.4 *±* 23.78 s for LEMON and 521.9 *±* 59.77 s for Cam-CAN, with an average data length ratio between the two resulting in 0.9128. Given the data length roughly matched and to retain as much MEG data as possible, we did not further shorten the Cam-CAN dataset to match LEMON. All data analyses delineated below took place in source space.

### 2.3 Hidden Markov Model

Once the data were source reconstructed, the next step is to identify dynamic RSNs and pinpoint the time periods during which they activate. The TDE-HMM is a generative model that can partition time series data into a sequence of recurring functional brain networks called *states* [20]. Each state represents an RSN characterised by distinct spatiotemporal patterns of spectral content. The basic structure of the TDE-HMM is depicted in Figure 1.

**Fig. 1.**
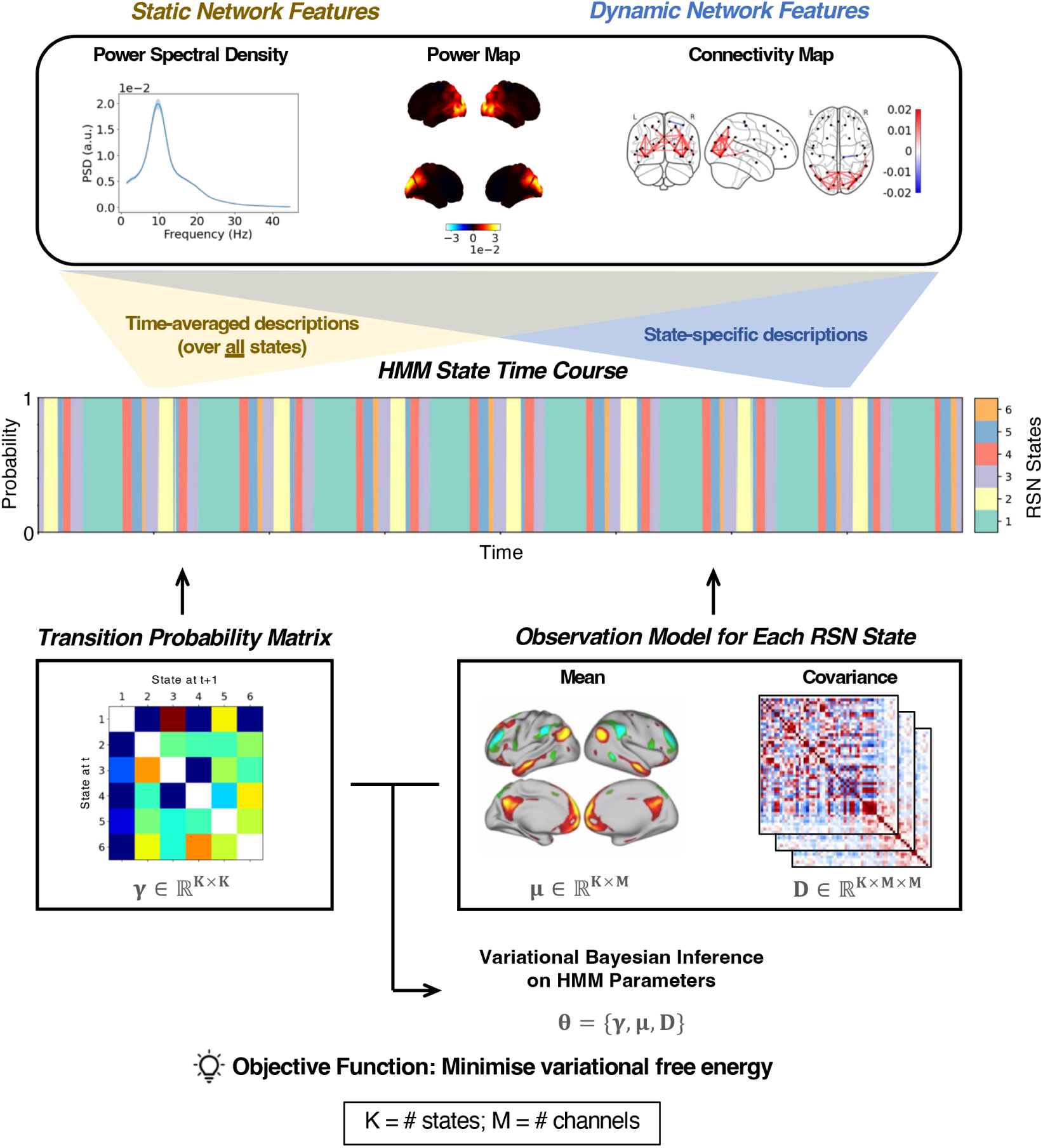
Overview of brain network feature analysis. This figure illustrates the methodology for extracting brain network features from MEG and EEG time series data. The brain network features we used consisted of power spectral density, spatial power distribution, and functional connectivity maps derived from spectral information within M/EEG recordings (top). The time series are segmented into discrete state activations via a Hidden Markov Model (HMM), resulting in an HMM state time course (middle). This process allows for the differentiation between dynamic network features—obtained from state-specific descriptions—and static network features, which are computed by time-averaging across all states. The HMM parameterises functional resting-state networks using a transition probability matrix and a multivariate Gaussian observation model with distinct means and covariances for each state by minimising variational free energy (bottom). The transition matrix quantifies the likelihood of transition from one state to another over consecutive time points, whereas the observation model elucidates the data generating process based on the latent (i.e., hidden state) representations. When combined with time-delay embedding, the HMM can identify states with distinct parcel power spectra and coherence networks (not shown).

The HMM typically comprises two primary components: a set of hidden states and an observation model for each state. Our observation model is a multivariate normal (Gaussian) distribution that has a distinct covariance matrix for each state. Crucially, the HMM is fit to TDE data, which augments the parcel time series with extra channels containing time-lagged versions of the original time series. Using TDE makes the state-specific covariance matrices sensitive to the power and frequency of oscillations in the original time series and results in states that have distinct power spectra for each parcel and coherence networks [23].

The HMM parameters were inferred using the variational Bayesian (VB) inference [32, 33]. This approach recasts the inference process as an optimisation problem, wherein the objective is to minimise the variational free energy. Using the VB method allows the model to learn uncertainty in parameter estimates as probabilistic distributions. To scale the learning process to large datasets, we utilised the stochastic version of the VB method adapted from [34]. As a result, the HMM outputs a time course of posterior probabilities, which signifies the probability of a state activation at each time point. By extracting the maximum a posteriori probabilities across all states by applying an argmax operation to these probabilities, we can generate a Viterbi path, or a *state time course*, delineating the most probable, yet mutually exclusive, state at each time point. The training process is described below. For more details on the design and implementation of the TDE-HMM, readers should refer to [20] and [23].

#### 2.3.1 Data Preparation

Before training a TDE-HMM model, the source reconstructed data were prepared to allow the model to learn states with unique spatio-spectral patterns. First, the data were time-delay embedded with *±*7 lags. With a 38-region parcellation, this embedding resulted in a total of 570 channels. PCA was then applied to reduce the dimensionality down to 80 channels to reduce memory requirements, and the transformed data were subsequently standardised over time. Since we are primarily interested in the transient patterns of spectral events, which are reflected in the state covariance matrices, we fixed the state mean vectors of the observation model to zero during the training process [23]. These prepared data served as the input training data, which was then shuffled and batched prior to model training.

#### 2.3.2 Hyperparameters

Training the model involves several key hyperparameters. First, the number of states was set to 6 to facilitate a reasonably low-dimensional data representation, which tend to be more reproducible and interpretable (see Section 2.3.5 for its validation). Next, we selected 64 as the batch size necessary for the stochastic VB inference. The input sequence length for batching the time series data was 800 samples. This length ensures minimal discontinuities between batched data, leading to less noisy updates of the parameter estimates. Lastly, the Adam optimiser [35] was used for updating the trainable parameters with a learning rate of 0.01 over a total of 20 training epochs.

#### 2.3.3 Model Initialisation

As the optimisation process for the HMM is stochastic, a certain level of variability in the final model parameters, referred to as *run-to-run variability*, is inevitable. To mitigate this, we use a number of random starts, or ‘initialisations’. This initialisation process was repeated five times, trained for two epochs in each repetition. The initialisation that yielded the lowest free energy was selected for the full model training. This approach has previously been shown to produce reproducible results [23].

#### 2.3.4 Model Runs

Despite the initialisation technique above, different model runs may converge to different local optima due to the stochastic nature of the parameter optimisation process. To obtain the best possible parameter estimates, therefore, the HMM model was trained ten times using the identical training dataset for both LEMON and Cam-CAN. From these 10 runs, a model with the lowest free energy was selected as the best run. This procedure was again repeated five times^3^, yielding a total of 50 runs (and 5 best runs) for each dataset. The best of the 5 best runs, hereinafter denoted as the *optimal* run, was chosen to be used for all subsequent analyses. As the order of inferred states is random for each model, the EEG and MEG states from their respective optimal HMM runs were manually aligned by eye^4^ based on their power and FC spatial patterns.

#### 2.3.5 Model Validation

Before the datasets were analysed, we validated that 6 states produce interpretable RSNs without any loss in training quality. For validation purposes, we trained 50 HMM runs with 8 and 10 states, following the method outlined in Section 2.3.4. For each set of dynamic RSNs (for 6, 8, and 10 states), we counted how many times a particular RSN appears across the best runs by qualitatively comparing its power spectral densities (PSDs), power maps, and FC networks (see Section 2.4). Results indicated that with 6 states, the HMM consistently converged to the same 6 RSNs across the best runs. In contrast, for 8 and 10 states, approximately 6 RSNs demonstrated consistency across the best runs, while the remaining states exhibited greater variability across runs for both EEG and MEG (i.e., the additional states corresponded to less reproducible RSNs). Figure A7 shows 10-state dynamic M/EEG RSNs, inferred from the optimal model run, as an example. The configuration of 6 states that guarantees the network features with the lowest model run-to-run variability was selected to ensure easier comparison between the modalities.

### 2.4 Network Feature Analysis

To compare the RSNs of the two modalities, we had to begin by extracting their static and dynamic network features. These features included PSDs, power maps, and FC networks, which were obtained by either averaging over the entire recording duration or computing state-specific descriptions based on the state time courses (Fig. 1). These features were qualitatively compared across modalities and evaluated for their efficacy in capturing age-related effects at the group level. Additionally, for the age classification task (see Section 2.7), we specifically computed TDE-PCA covariance matrices and summary statistics as input network features. The network feature computations outlined in this subsection were largely adopted from [5] and conducted using the OSL Dynamics toolbox [23].

When interpreting these network features, it is important to note that the approach we used for source reconstruction (i.e., unit noise gain invariance LCMV beamformer) implicates the loss of absolute amplitude information from the sensor recordings. This step was necessary to address the depth bias in source reconstruction, but it precludes us from making any direct comparison of absolute network features across modalities. Furthermore, we standardised (i.e., z-transformed) the data across subjects before computing network features (see below), which eliminates amplitude information across individuals. This procedure allows us to compare the relative network features in each frequency band across modalities and subjects. Therefore, all network features employed in this paper are measures relative to the time-average for each sensor and subject, not absolute measures.

#### 2.4.1 General Linear Model

We employed a group-level general linear model (GLM) to study age-related effects and measure the differences in various network features between the young and old groups. The value of a network feature (e.g., power for a parcel, or FC for a parcel-pair, at a particular frequency) as it varied of subjects was fed in as the dependent variable to the group GLM, and the GLM was fitted independently to each network feature. The group-level GLM design matrix (Fig. A2) contained two regressors that modelled the group means of the young and old cohorts separately. Sex and head size were included as covariates to remove any confounding effects introduced by these variables. In particular, the GLM contained two sex regressors, one for male and the other for female, and one head size regressor, all of which were demeaned prior to model fitting. From the fitted GLMs we computed the mean group differences as a contrast of parameter estimates (COPEs). Group-averaged values of each network feature were simply taken from the regression estimates of the fitted GLMs. For a comprehensive overview of the GLM and its application to neuroimaging data, readers should consult [37].

#### 2.4.2 Power Spectral Densities

First, we computed subject-level PSDs to examine the distribution of power across distinct frequency ranges. For the static analysis, we employed Welch’s method [38] on the original parcel time course to calculate static PSDs of size [*N × M × P*], where *N* is the number of subjects, *M* the number of parcels, and *P* the number of frequency bins. Each PSD was calculated using a 2-second Hann window with 50% overlap across a 1.5-45 Hz frequency range, and each parcel time course was standardised (to have zero mean and unit variance) at the individual subject level before calculation. To compare the static PSDs between the young and old cohorts, PSDs were fitted to a GLM, producing the group-averaged PSDs of size [*M × P*].

For the dynamic analysis, the power spectrum for each subject, region, and state was separately calculated using the multi-taper method, which is the typical approach used with TDE-HMMs [39]. As in the static analysis, the data were standardised at the individual subject level. PSDs for each state were estimated using the source data and inferred state time course, which helped identifying time points corresponding to the active states. For each subject and state, a multi-taper spectrogram was generated over the entire time points during which a single state was activated, using 7 tapers with a time-half bandwidth of 4. By averaging a spectrogram over time, state-specific multi-taper spectra (i.e., dynamic PSDs) of size [*N ×M ×P*] were obtained. In addition, state-specific coherence spectra of size [*N × M × M × P*] were calculated using the multi-taper cross-spectral densities. For visualisation purposes, the mean PSD across all states, weighted by the duration of state activations, was subtracted from the state-specific multi-taper spectra at the subject level, and the group-averaged dynamic PSDs were computed by averaging over the subjects in each group.

#### 2.4.3 Power Maps

Power maps were generated by averaging PSDs across specified frequency ranges of 1.5 to 20 Hz (wide-band), 1.5 to 4 Hz (delta), 4 to 8 Hz (theta), 8 to 13 Hz (alpha), and 13 to 20 Hz (beta). These frequency bands were manually defined, rather than by using the non-negative matrix factorisation method employed in [23], due to the failure of EEG data to converge during factorisation. For static analyses, power maps were derived from static PSDs; for dynamic analyses, we used dynamic, state-specific multi-taper spectra. Mean group differences in static power maps were quantified as COPEs by fitting the power maps to GLMs. For visualisation purposes, power values across brain regions were voxel-weighted and interpolated. No threshold was applied to the plots. In dynamic power maps, the mean power map across all states, weighted by the duration of state activations, was subtracted from the state-specific power maps at the subject level.

#### 2.4.4 FC Networks

FC was measured as the amplitude envelope correlation (AEC) for static analyses and coherence for dynamic analyses. The preference for AEC in static analyses was due to the method’s known consistency in capturing stationary connectivity measures with relatively high test-retest reliability [40]. Amplitude envelopes were computed by applying the Hilbert transform to the source data, which were band-pass filtered between 1.5 to 20 Hz and subsequently standardised. The static AEC networks were estimated by taking pairwise correlations of these amplitude envelopes across brain regions for each subject.

Conversely, the dynamic coherence networks were derived by averaging a coherence matrix for each state across 1.5-20 Hz. Note that it is conventional to use AEC for the static analysis and coherence for the dynamic analysis. Since the HMM infers coherence networks given the time-delay embedded data, using coherence is better suited to the dynamic case [23]. Moreover, applying AEC to dynamic FC usually involves segmenting signals according to state time courses and concatenating these segments. This process, however, can introduce edge effects and phase discontinuities, which may be detrimental to the Hilbert transform and its amplitude envelope. On the other hand, we opted not to use coherence for static analyses because it is reported to perform poorly in static connectivity estimation, showing low test-retest reliability [40].

A threshold corresponding to the 95th percentile was applied to the FC networks for visualisation. In dynamic analyses, FC networks averaged across all states, weighted by the duration of state activations, were subtracted from the state-specific FC maps.

#### 2.4.5 Summary Statistics of Network Dynamics

In the age group classification task (outlined in Section 2.7), we incorporated summary statistics detailing the dynamics of RSNs as part of the input features for our classifier. Four summary metrics were computed to quantify the temporal characteristics of the HMM states for each subject [4, 23]. These metrics were calculated using the state time courses and include:

- **Fractional occupancy**: the proportion of the total time spent in each state.
- **Mean lifetime**: the average amount of time spent in each state before transitioning into another state.
- **Mean interval**: the average amount of time elapsed between visits to the same state.
- **Switching rate**: the number of state visits per second.

#### 2.4.6 TDE-PCA Covariance Matrix

Another input feature we used for the classification task was static and dynamic TDE-PCA covariance matrices. These features summarise the spatiotemporal properties of the source data.

Subject-specific static TDE-PCA covariance matrices were generated by calculating the covariance of the TDE parcel data for each subject, performing a PCA to reduce the data to a rank of 80 and then standardising the resulting PCs. Note that this corresponds precisely to the steps taken in the data preparation to compute what we refer to as the standardised PCA-reduced TDE data (c.f., Section 2.3.1).

The HMM explicitly infers dynamic TDE-PCA covariance matrices. However, these values are group-level estimates. To calculate subject-specific estimates of the dynamic TDE-PCA covariance, we used the dual-estimation method [41, 42], where we fixed the state probabilities in the group-level HMM and re-estimated the state TDE-PCA covariance values using the subject-wise prepared source data.

### 2.5 Statistical Analysis

Having extracted the static and dynamic RSN features, we next examined the statistical differences between the two age groups to probe how age-related effects are represented by each modality. We employed non-parametric permutation tests for all pertinent brain network features, using t-statistics as the permuted statistic [43, 44]. For PSDs, we determined group-level significance using a two-tailed cluster permutation test. Clusters were formed over frequencies, with a cluster-forming threshold set to 3. For power maps, we utilised a two-tailed max-t permutation test to evaluate group-level significance.

For both types of permutation tests, a metric of interest was first fitted to a group-level GLM as previously described. From the fitted GLMs, we computed COPEs (i.e., mean group differences) and their variances, from which t-statistics could be calculated [37]. The group regressors were permuted 5,000 and 10,000 times to construct a null distribution for the cluster and max-t permutation tests, respectively. We then selected a significance threshold from this null distribution for statistical testing.

Due to the multiplicity of different non-parametric permutation tests, we adjusted for multiple comparisons across frequency bands and states using Bonferroni correction. For the number of bands and states we accounted for, please refer to the figure legends for further details. This correction was applied within the MEG or EEG dataset; tests repeating across the modalities were not treated as part of the same statistical family.

### 2.6 Reproducibility of Static and Dynamic RSNs

To gauge whether the static RSNs derived from the EEG and MEG data are reproducible, we split each dataset into two halves and inferred networks on each half separately [5]. For each half, we compared static PSDs, power maps, and FC networks both within and across modalities. PSDs were averaged across subjects and parcels, while power maps and FC networks were averaged over a frequency range of 1.5-20 Hz and across subjects.

In Section 2.3.4, it was noted that the HMM is prone to converging to local optima, resulting in the model potentially arriving at similar variational free energies with different parameter estimates. Therefore, in addition to the initialisation method and the selection of the optimal runs, we implemented an additional validation step that assessed reproducibility across split-half datasets. Specifically, the LEMON and Cam-CAN datasets were each divided into two halves, with separate HMMs trained on each subset. For each half, a model was trained ten times.

We assessed the reproducibility of the network dynamics through three measures. First, we examined the spatial distribution of power within and across the modalities by comparing the state-specific power maps derived from each of the split-half data. Next, we quantified the similarity of group-level state covariance matrices both within and across modalities by computing RV coefficients for pairs of matrices [45]. Lastly, we evaluated the consistency of inferred transition probability matrices within and across modalities using the Jensen-Shannon (JS) distance [46]. This metric was chosen for its symmetric nature and finite values, making it suitable for comparing probabilistic distributions. The JS distance was calculated for each row of transition probability matrices, and these values were averaged over all rows. This approach treats each row as a distinct probability distribution, viewing the transitions from one state to every other as separate entities and considering the overall similarity across these distributions. It should be noted that power maps and RV coefficients were calculated using the optimal HMM run with the lowest variational free energy. The JS distances, on the other hand, were computed using all 10 runs. For each comparison within and across modalities, a distribution of JS distances was obtained from permuted pairs of 10 HMM runs across two split-half datasets. We additionally tested whether the JS distances within EEG and MEG are drawn from the same distribution using the Wilcoxon rank-sum test, after checking the assumptions of normality and homogeneity of variances.

### 2.7 Age Group Classifications

Finally, to evaluate the degree to which the network descriptions derived from MEG and EEG are meaningful, we assessed their utility in predicting age groups by employing a logistic regression classifier with L2 regularisation [47, 48]. The task incorporated three distinct categories of input features: static, dynamic, and a combination of both. Static features comprised static TDE-PCA covariances, while dynamic features included dynamic TDE-PCA covariances alongside summary statistics representing network dynamics. These input features were concatenated before training a classifier.

Separate classifiers were trained for each category of input features with a nested cross-validation approach, which involved two levels of cross-validation. In this framework, the inner cross-validation focused on hyperparameter optimisation, while the outer cross-validation estimated the generalisation error across multiple test sets. In the outer loop, input features were randomly shuffled and subjected to a 5-fold cross-validation. The training set partitioned from this outer loop underwent another 5-fold cross-validation within the inner loop. For the inner cross-validation, we standardised the data and applied dimensionality reduction (i.e., PCA with whitening) before fitting any classifiers to enhance predictive accuracy. The optimal number of principal components (PCs) was determined via a grid search on the inner training folds over 10, 20, and 30 PCs.

The best-performing classifier from the inner loop was then selected and evaluated on the test set in the outer loop to compute its overall predictive accuracy. Here, the test set was processed using the same standardisation and PCA parameters fitted on the training set to ensure consistency across data splits and maintain fair model evaluation. This entire nested cross-validation procedure was repeated 10 times. We calculated and reported the mean and standard deviation of the accumulated test accuracies to provide a comprehensive overview of the classifier’s performance on the M/EEG network features.

To determine if the test accuracies exceeded the level anticipated by random chance alone (i.e., 50%), a one-tailed, one-sample t-test was used. We employed a two-tailed, two-samples t-test to assess whether there are significant differences in the prediction scores between MEG and EEG, as well as to examine if the prediction scores varied depending on how the data were reconstructed (i.e., the use of subject sMRI data in source reconstruction). The significance of all tests was evaluated against a Bonferroni-corrected threshold, adjusted for three categories of input features. The assumptions of normality and homogeneity of variances were verified prior to conducting these tests. When the assumption of equal variances was violated, we employed Welch’s t-test as an alternative to the standard two-sample t-test.

### 2.8 Use of Artificial Intelligence Assistance

While preparing this paper, the authors used ChatGPT to refine the styles and grammar of the parts of the written text. This tool was never used to create and write new content. The authors reviewed and edited any changes made and take full responsibility for the content of the publication.

### 2.9 Data and Code Accessibility

For the original LEMON and Cam-CAN datasets, the readers should refer to [21] and [22], respectively. All codes for analyses and data visualisations, including scripts for preprocessing, source reconstruction, and model training, were written in Python and shared at https://github.com/OHBA-analysis/Cho2024 MEEG RSN. Some of the scripts were built upon the OSL Dynamics software [23]. The inferred HMM parameters and computed RSN features are shared in the same repository. Further requests for prepared data or concerns regarding the codes should be directed to and will be fulfilled by the authors.

## 3 Results

### 3.1 Static Analysis

#### 3.1.1 Static RSN features are qualitatively similar across the modalities

As discussed in the Introduction, the comparability between static functional brain network features of MEG and EEG has been established by previous literature. To ascertain this similarity with a medium-density EEG (61 channels), we examined how static RSN features qualitatively compare between these modalities (Fig. 2). The subject-averaged, wide-band (1.5-20 Hz) power maps for EEG and MEG both revealed pronounced activation in the frontal, parieto-occipital, and sensorimotor cortices. MEG exhibited a higher signal-to-noise ratio (SNR), with its power values higher than EEG. The subject-averaged, wide-band FC maps also indicated similarities between the modalities, in which the connections were strongest at the parieto-occipital region. Here, EEG showed stronger connection strengths, whereas MEG FC was more concentrated on the parietal cortex. Likewise, M/EEG PSDs averaged across subjects and parcels displayed comparable spectral distributions with a prominent unimodal peak in the alpha frequency band. However, their aperiodic activities diverged, as EEG showed a steeper decay following a 1/f pattern. All these trends were also evident in the M/EEG data that had been reconstructed using a standard brain structural image instead of subject-specific sMRI images (Fig. A3).

**Fig. 2.**
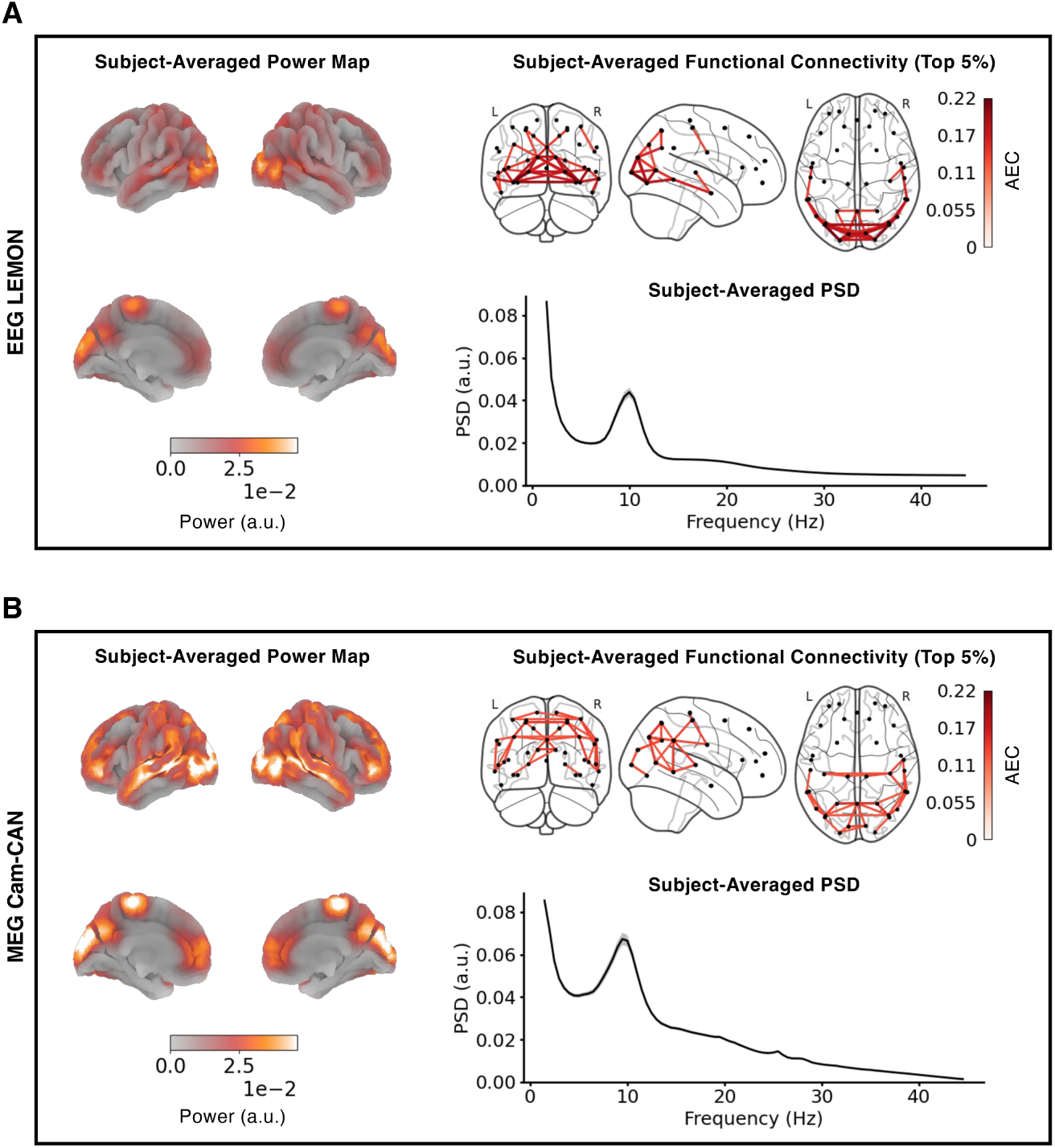
Qualitative comparison of static resting state networks between EEG and MEG. **(A)** Brain surface map of static power for each brain parcel, calculated by first averaging PSDs across the frequency range of 1.5-20 Hz at the individual level and then across all subjects (left). On the top right, an FC network of static AEC averaged across subjects, with only the top 5% of correlation strengths shown for better visualisation. AEC values were calculated on envelopes obtained from subject-wise parcel time courses, band-pass filtered from 1.5-20 Hz. On the bottom right, the static PSD averaged across both brain parcels and subjects, with standard errors across parcels indicated by grey shading. **(B)** The subject-averaged static power map, functional connectivity network, and PSD generated using the MEG data following the same analyses as in (A).

To ascertain whether the static RSNs derived from M/EEG and their overall comparability are reproducible, we extracted the same static PSDs, power maps, and FC networks from the EEG and MEG datasets divided into halves. We then assessed their reproducibility within and across the modalities (Fig. 3). A qualitative comparison illustrated that static spectral, power, and connectivity distributions remained consistent within each modality. In both EEG and MEG, each half highlighted identical brain regions with high power and connectivity values, while also demonstrating similar PSD shapes (Fig. 3A, B). Across the modalities, any comparisons of two halves mirrored the observations from the full dataset. That is, MEG showed higher power than EEG, whereas EEG exhibited stronger FC than MEG, with MEG accentuating the parietal cortex relatively more. To this extent, the comparability of the static RSN features was higher within each modality than between the two modalities.

**Fig. 3.**
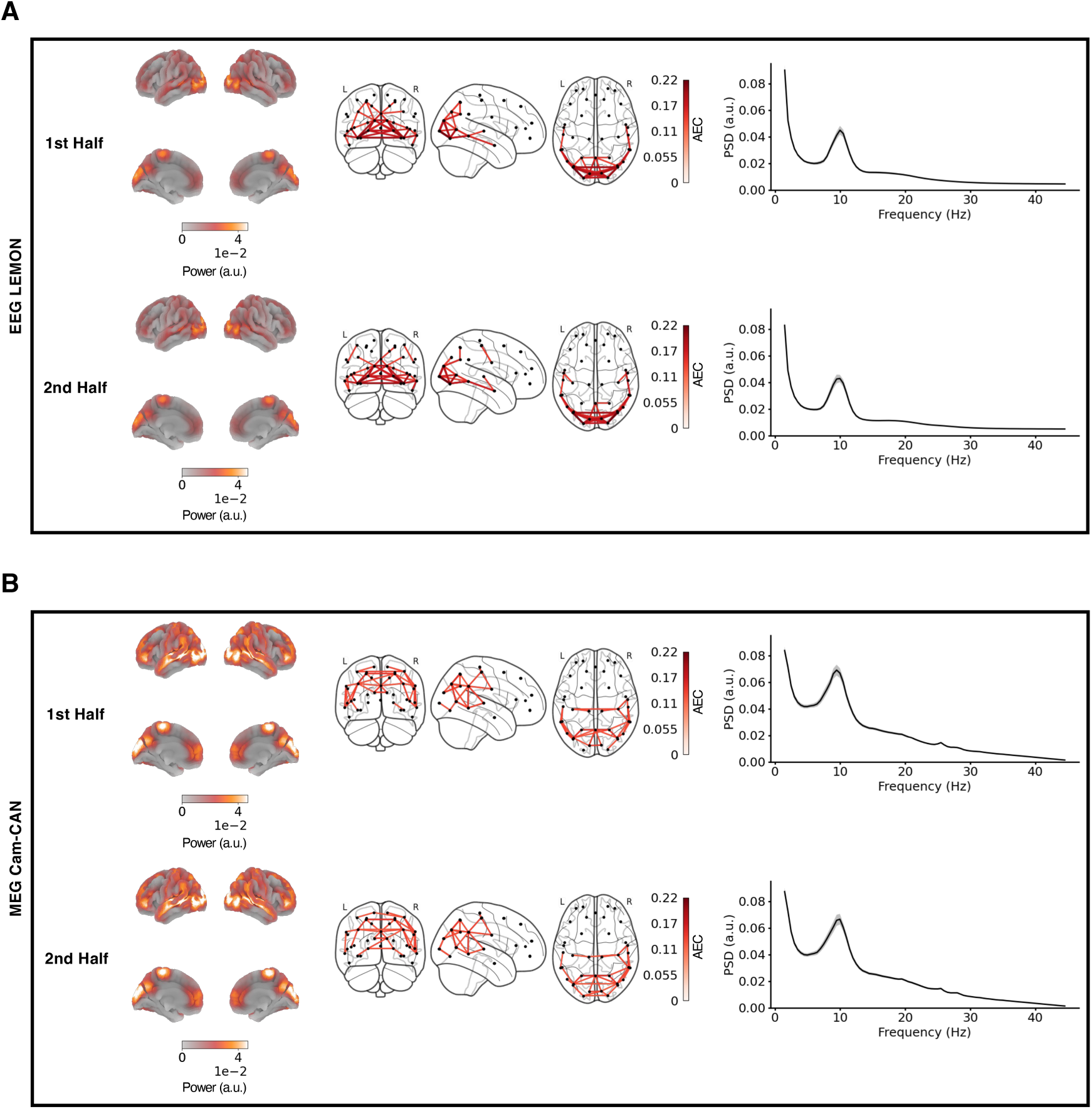
Resting-state network reproducibility of static network features within and across modalities. **(A)** The subject-averaged, wide-band (1.5-20 Hz) power maps and FC networks, along with the PSDs averaged across both brain parcels and subjects, computed from the first half (top panel) and the second half of the resting-state EEG data (bottom panel). Each half comprises data from 48 subjects. The power maps depict lateral surfaces at the top and medial surfaces at the bottom, and the FC networks show only the top 5% of correlation strengths. For the PSDs, standard errors across parcels are indicated by grey shading. **(B)** The same analysis as in (A) is conducted using the MEG data.

In summary, EEG and MEG demonstrated qualitative correspondence across all static network features, despite some discrepancies. This comparability was noted in the data that were reconstructed using a standard brain structural image as well, with which we could observe the same discrepancies between the two modalities. These network features also displayed a good level of reproducibility, both within and across the modalities, when qualitatively contrasted.

#### 3.1.2 MEG and EEG reveal comparable age effects in static network features

To investigate whether MEG and EEG provide equally meaningful static network descriptions, we examined age-related effects captured by their static RSN features. Both modalities demonstrated comparable age-related effects in PSDs, identifying statistically significant differences between the age groups across the delta, theta, and beta bands (Fig. 4A, F). However, slight discrepancies were noted at higher frequencies (30-45 Hz). The topographic maps highlight the alpha peak PSD differences, the power differences in lower frequencies, and the power differences in the beta band; these topographical patterns additionally showed qualitative similarities between the two modalities.

**Fig. 4.**
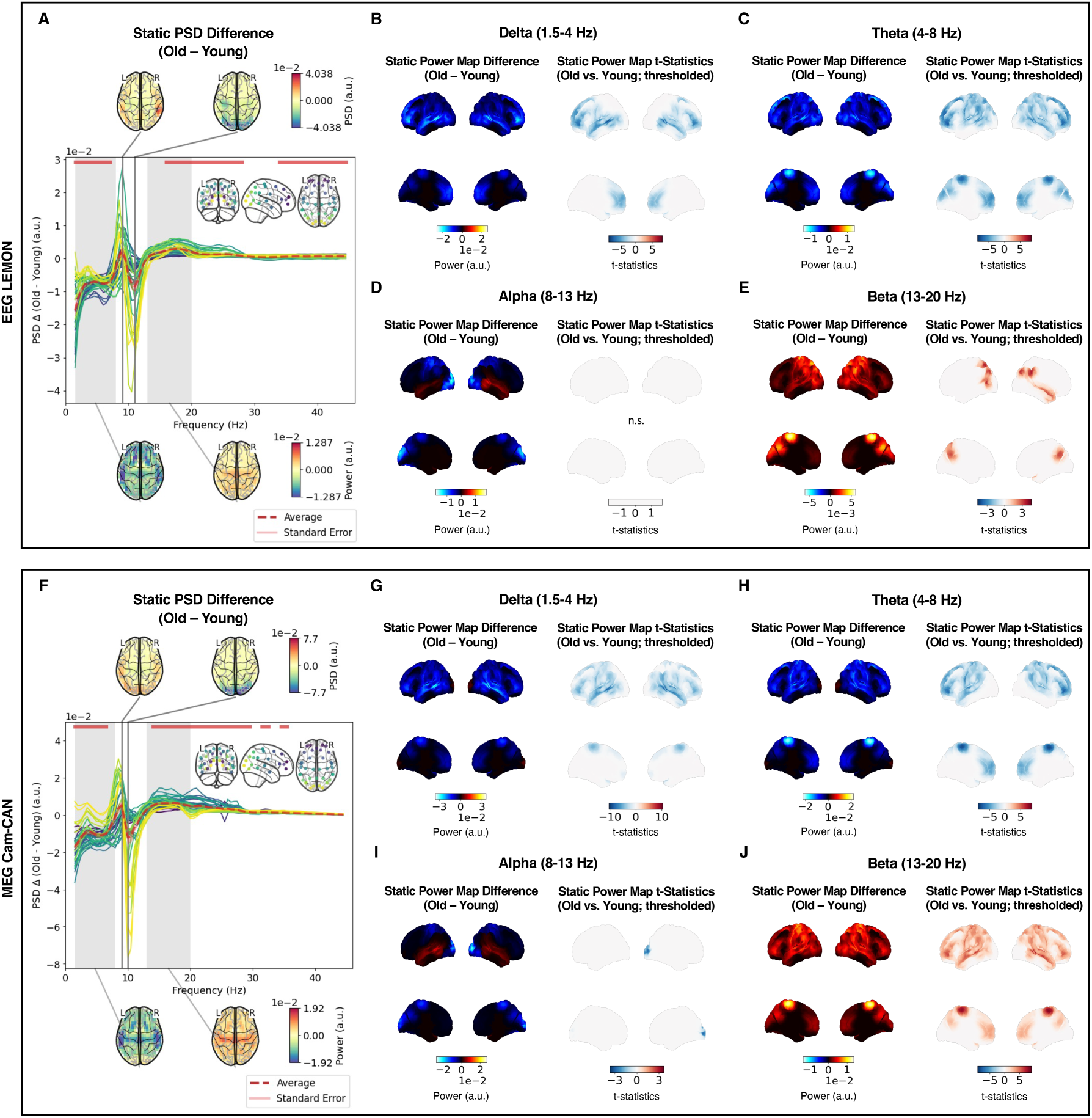
EEG and MEG reveal comparable age-related effects in static network features. **(A)** The between-group PSD difference (red dotted line), averaged over parcels and subjects, is depicted alongside green lines denoting the subject-averaged PSD differences for each parcel (the inset in the top-right corner indicates the shade of green used for each brain parcel) of the EEG data. Frequency ranges with significant PSD differences (p *<* 0.05; horizontal red bars) are identified via a cluster permutation test on parcel-averaged PSDs of the two age groups. Additionally, the top topographical maps show the peak/trough group-level PSD differences found in the alpha frequency band. The bottom topographies show group-level PSD differences integrated separately across the lower (1.5-8 Hz) and beta (13-20 Hz) frequency bands, represented by the grey shading. **(B)** On the left, a brain surface map illustrating delta band power differences between the old and young groups is plotted for the EEG data, with lateral surfaces at the top and medial surfaces at the bottom. On the right, the max-t permutation test (p *<* 0.05; Bonferroni-corrected, n=4 frequency bands) was applied to the power map to highlight the age-related effects, with significant parcels accentuated. Red and blue colours indicate higher power in old and young participants, respectively, and non-significant findings are labelled as n.s. The same analysis as in (B) is shown for the **(C)** theta, **(D)** alpha, and **(E)** beta band EEG power maps. **(F)** The same analysis as described in (A) is shown for the MEG data. **(G)-(J)** The same analysis as described in (B) is shown for the delta, theta, alpha, and beta band power differences, respectively, using the MEG data.

Further analysis of age-related effects in narrow-band (delta, theta, alpha, beta) power spatial maps indicated that the general trend of power differences between the age groups (whether young participants exhibited higher power than old participants, or vice versa) was statistically significant across two modalities (Fig. 4B-E, G-J; right). Nonetheless, their specific spatial distributions still demonstrated slight discrepancies. In the delta power map, the young cohort exhibited higher power in the frontal and temporal cortices, and specifically in the sensorimotor region in MEG (Fig. 4B, G). In the theta power map, the young group showed increased power in the frontal and sensorimotor regions, with only EEG displaying higher power in the parietal region (Fig. 4C, H). Significant age-related alpha effects were exclusively observed in MEG within the occipital cortex, whereas EEG did not identify any regions with notable age effects (Fig. 4D, I). In the beta power map, age-related effects were more prominent in MEG, which identified effects in the parietal, frontal, and sensorimotor cortices, albeit both modalities recognised effects in the parietal region (Fig. 4E, J). The general trend of between-group static power differences was more evident in the qualitative representation of age effects (Fig. 4B-E, G-J; left).

The observed age-related effects remained consistent in the M/EEG data that were reconstructed using a standard brain structural image (Fig. A4). However, notable differences were identified, including minor deviations in spatial patterns within the parietal and temporal cortical regions in the EEG beta power map (Fig. A4E), along with the absence of age-related effects in the MEG alpha power map (Fig. A4I).

In summary, although some of the spatial details of the age-related effects varied, MEG and EEG often captured group-level differences of a similar nature. Their comparability persisted even when the data were reconstructed with a standard brain structural image. Nonetheless, there was a tendency for MEG to be slightly more sensitive to age-related changes.

### 3.2 Dynamic Analysis

#### 3.2.1 Dynamic RSN features are qualitatively similar across MEG and EEG

Following the static analysis, we examined how dynamic RSN features compared between the two modalities. Figure 5 presents the subject-averaged PSDs and wide-band (1.5-20 Hz) power map and FC network for each state, demonstrating their alignment between EEG and MEG. Notably, the spatial patterns of power distributions showed a marked resemblance between the modalities. Nonetheless, distinctions arose particularly in states 2 (the visual network) and 6 (the sensorimotor network), where there was relatively less correspondence. These states exhibited divergent areas of stronger activation or deactivation between the modalities; for instance, state 2 displayed elevated occipital activity in EEG, while more pronounced deactivation in non-occipital areas was observed with MEG.

**Fig. 5.**
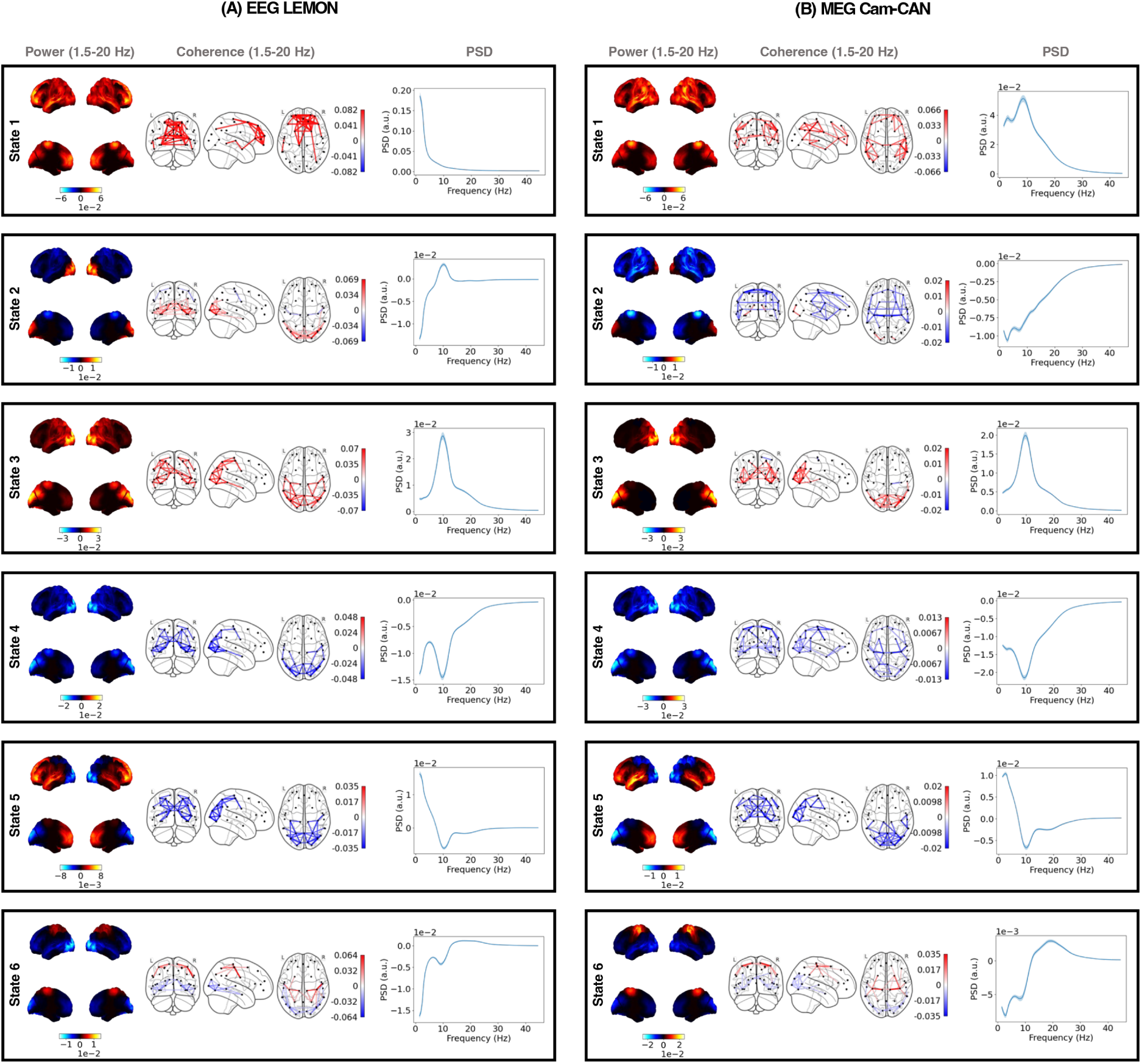
Qualitative comparison of dynamic resting state networks between EEG and MEG. **(A)** Each box shows the subject-averaged power map (left), FC network (middle), and parcel-averaged PSD (right) of each HMM state for 96 EEG subjects. Each state is considered as a distinct RSN. The power maps show lateral surfaces at the top and medial surfaces at the bottom. The FC networks illustrate connections with the top 5% coherence values (irrespective of sign). The shaded areas of the PSDs represent the standard error of the mean across the parcels. The power maps, FC networks, and PSDs are visualised relative to their average across all states. **(B)** The plots follow the same format as (A), showing the RSNs computed from 96 MEG subjects.

Similarly, the state-specific FC networks and PSDs largely maintained a high degree of comparability between EEG and MEG, alongside more detailed discrepancies. Such discrepancies were most pronounced in states 1 (a putative anterior default mode network) and 2 (the visual network), in which the top 5% coherence values and PSDs highlighted different brain regions and spectral patterns, respectively, when visualised relative to the average values across all states.

The comparability of dynamic network features across the two modalities was reflected in the MEG and EEG RSNs derived from the data reconstructed using a standard brain structural image (Fig. A5). Importantly, the discrepancies previously observed persisted, with states 2 and 6 displaying reduced correspondence in power maps and states 1 and 2 showing lesser alignment in FC networks and PSDs.

A possible source of the between-modality discrepancies might stem from variability among subjects and between different model runs. To ensure that the overall consistency of the dynamic RSNs derived from EEG and MEG are replicable despite these minor differences, we evaluated the reproducibility of HMM states within and across the modalities (Fig. 6). We started by comparing the state-specific power maps generated from the EEG and MEG datasets split into two halves. Our qualitative comparison revealed that spatial power distributions remained consistent within each modality (Fig. 6A, B). There was also a notable similarity in power maps across the modalities, with state 6 being the sole exception to this trend.

**Fig. 6.**
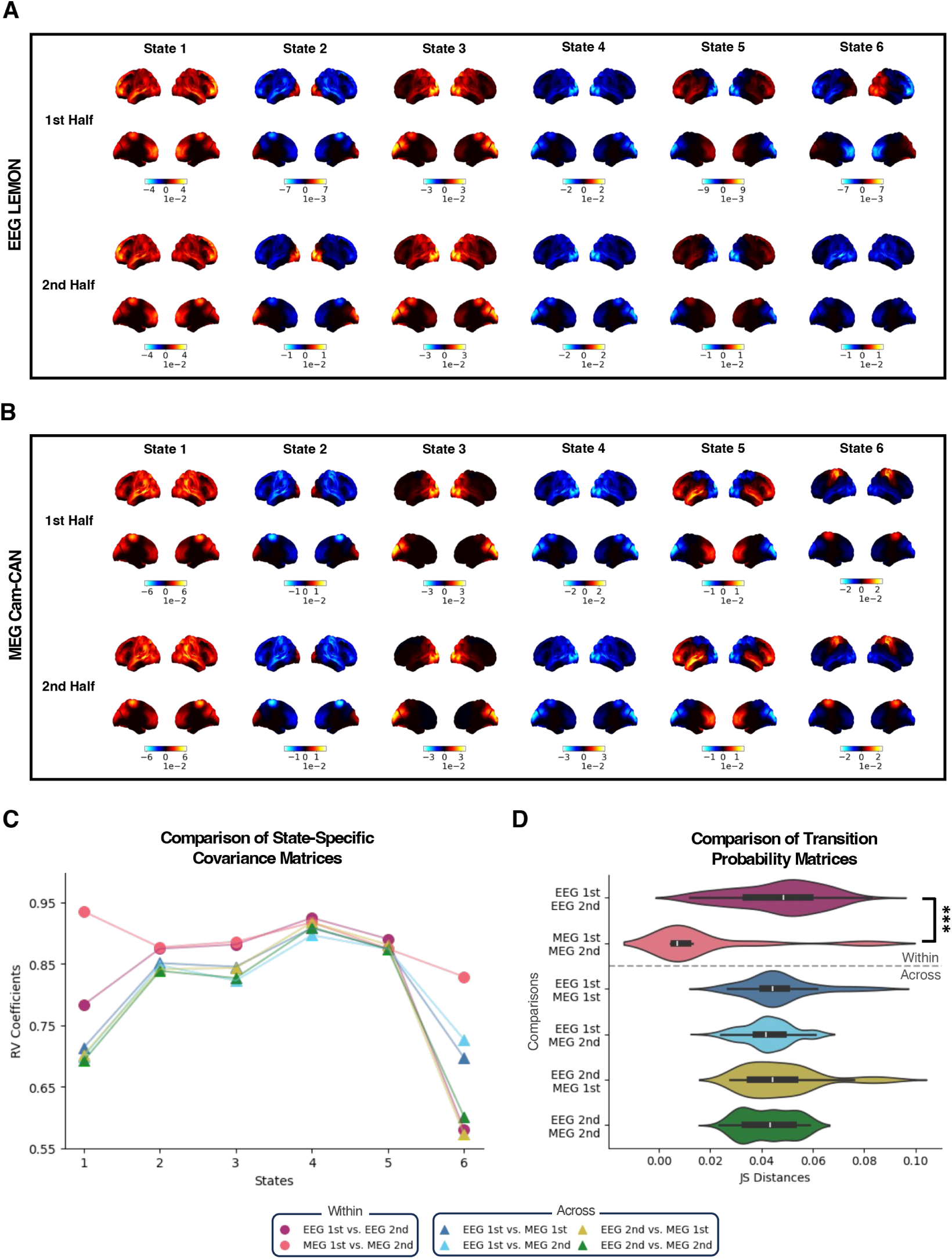
Resting-state network reproducibility of dynamic network features within and across modalities. **(A)** The wide-band (1.5-20 Hz) power maps for resting state EEG data, averaged across subjects, are shown for the first half of the dataset (top) and the second half of the dataset (bottom), with each half comprising data from 48 subjects. These maps show lateral surfaces at the top and medial surfaces at the bottom, visualised relative to their average across all states. **(B)** The same analysis as in (A) is conducted using the MEG data. **(C)** For each state, the similarities between inferred covariance matrices within and across modalities are measured as RV coefficients (higher value indicates more reproducible). **(D)** Similarities between inferred transition probability matrices within and across modalities are measured as the Jensen-Shannon (JS) distances and depicted in bar graphs (lower value indicates more reproducible). The statistical difference between JS distances within EEG and MEG was tested using the Wilcoxon rank-sum test (*U* =9.053, *p*=1.394e-19). Significance levels are indicated by ***: p *<* 0.05, **: p *<* 0.01, *: p *<* 0.001.

The reproducibility of HMM states was also measured with respect to the inferred HMM parameters. In particular, we computed the RV coefficients, a generalisation of the squared Pearson correlation coefficient, between the covariance matrices of two split-half M/EEG datasets (Fig. 6C). As anticipated, RV coefficients for comparisons within the same modality were typically higher (more reproducible) than those for comparisons across the modalities. For all states except state 6, RV coefficients exceeded 0.65 for any given comparison, although the variability of this measure over different types of comparisons was relatively higher for states 1 (a putative anterior default mode network) and 6 (the sensorimotor network). Notably, RVs within MEG were generally higher than those within EEG.

Subsequently, we quantified the difference between transition probability matrices of two split-half datasets as the JS distance (Fig. 6D). This distance was lower (more reproducible) for the comparison within MEG than that within EEG or across the modalities. In particular, the JS distances within MEG were significantly smaller than those within EEG. The comparisons across the modalities had median JS distances similar to the median value within EEG, although the latter exhibited a larger variance, including relatively lower distances. Such similarity partly serves as additional evidence that EEG and MEG RSNs are comparable as they pertain to inferred state transition probabilities.

In summary, EEG and MEG were broadly similar across the dynamic network features, while also showing some discrepancies. Their comparability, as well as discrepancies, persisted when the data were reconstructed using a standard brain structural image. Furthermore, we measured the reproducibility of these network features, as well as of the HMM parameters, both within and across the modalities. This showed that state 6 (the sensorimotor network) was the least reproducible and that EEG was less reproducible than MEG.

#### 3.2.2 MEG and EEG reveal comparable age effects in dynamic network features

The age-related effects in dynamic PSDs and power maps derived from MEG and EEG are illustrated in Figure 7. Note that for each dynamic measure we compute the “dynamic contribution to the state-specific” measure, resulting in any dynamic effects above the static effects. For example, the power in a state in a particular frequency band is the power in that state in addition to the static power in that frequency band.

**Fig. 7.**
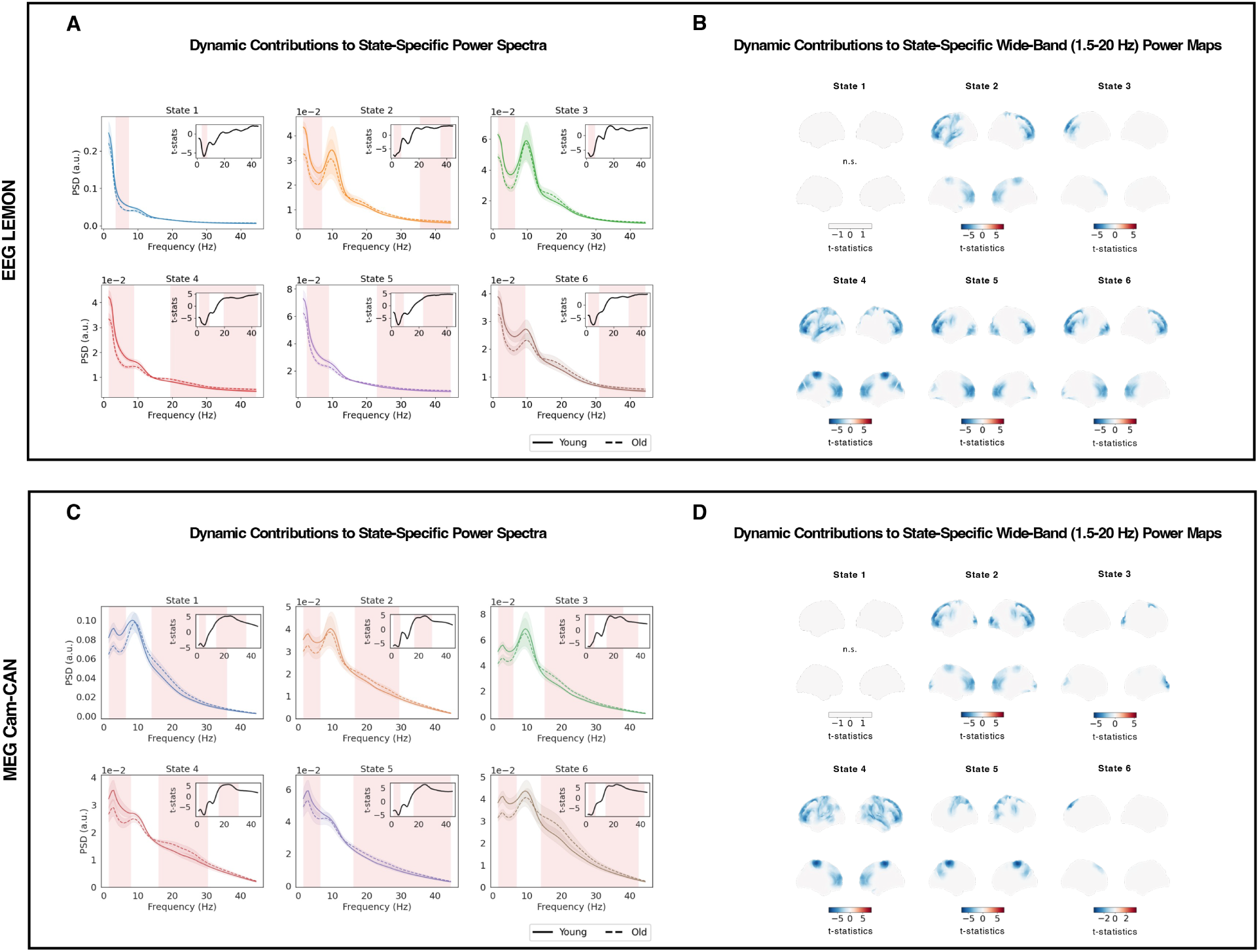
EEG and MEG reveal comparable age-related effects in dynamic network features. **(A)** The state-specific, dynamic PSDs of the EEG data averaged across grouped subjects and parcels, depicted for old (dotted) and young (solid) participants. For each HMM state, a cluster permutation test is conducted on parcel-averaged PSDs to detect age-related dynamic effects, with frequencies exhibiting statistical significance (p *<* 0.05; Bonferroni-corrected, n=6 states) highlighted in red. An inset at the top-right corner displays the t-statistics, signifying the magnitude of age-related effect sizes between the groups. **(B)** The state-specific power maps with significant between-group differences for the EEG data. These differences are validated through max-t permutation tests (p *<* 0.05; Bonferroni-corrected, n=6 states), with significant parcels accentuated. Red and blue colours indicate higher power in old and young participants, respectively, and non-significant findings are labelled as n.s. **(C)** The same analysis as in (A) performed on the MEG data, marking significant age-related dynamic effects in state-specific PSDs. **(D)** The same analysis as in (B) performed on the MEG data, illustrating significant age-related dynamic effects in state-specific wide-band power maps. Note that in all cases we plot the “dynamic contribution to the state-specific” feature, resulting in dynamic effects that are above the static effects.

For both modalities, the age-related effects for the dynamic PSDs (Fig. 7A, C) broadly appeared in the same frequency bands as for the static PSDs. Specifically, (Figure 7A, C) shows significant spectral differences between the age groups in the delta and theta bands across all states for both MEG and EEG. However, consistency in these differences was less evident in the beta band and higher frequencies. While states 4, 5, and 6 exhibited overlapping effects in these frequencies, states 1, 2, and 3 did not. In state-specific, wide-band (1.5-20 Hz) power maps, all states except state 1 displayed roughly equivalent age-related effects, with the young cohort showing higher power compared to the old group (Fig. 7B, D). However, states 2, 4, and 6 presented overlapping patterns across brain regions between the two modalities, whereas states 3 and 5 revealed significant power differences in regions that varied across the modalities. The observed age-related effects remained consistent in the M/EEG data that were reconstructed using a standard brain structural image without subject sMRI images, although to a lesser degree compared to the static analysis (Fig. A6). In particular, the age effects in higher frequencies altered in the PSDs of EEG states 2 and 4 (Fig. A6A). The differences were more pronounced on state-specific power maps. In general, significant power spatial distributions exhibited different patterns when a standard structural image was used. While EEG and MEG state 1 highlighted significant age-related effects in the occipital and parietal cortices, respectively, significance in MEG state 6 diminished under this condition (Fig. A6B, D).

In summary, MEG and EEG demonstrated broadly comparable age-related effects in dynamic network features (specifically, the dynamic effects above the static effects), albeit to a lesser extent compared to static features. This comparability persisted when the data were reconstructed using a standard brain structural image, although it was not maintained at a level similar to the static analysis.

### 3.3 Age Group Classification Task

Our comparative analysis of how MEG and EEG capture age-related effects in static and dynamic RSN features showed that both modalities offer broadly comparable network descriptions. To extend this analysis further in a quantitative manner, we aimed to estimate the predictive power of these network features in accurately classifying subjects into distinct age groups. Utilising three distinct sets of input features—static, dynamic, and a combination of both—we employed a logistic regression classifier with an L2 regularisation penalty (see Section 2.7).

We evaluated prediction scores against test datasets across 10 repeated classification tasks, with Figure 8 illustrating the resulting accuracies in age prediction. Both modalities showed predictions that exceeded what would be expected by random chance alone (i.e., 50%); the mean prediction accuracies, averaged across 10 repetitions, were significantly higher for MEG than EEG when employing static and dynamic input features (Fig. 8A). Although this statistical difference was not observed with combined features, there was still a clear trend towards better prediction for MEG compared to EEG.

**Fig. 8.**
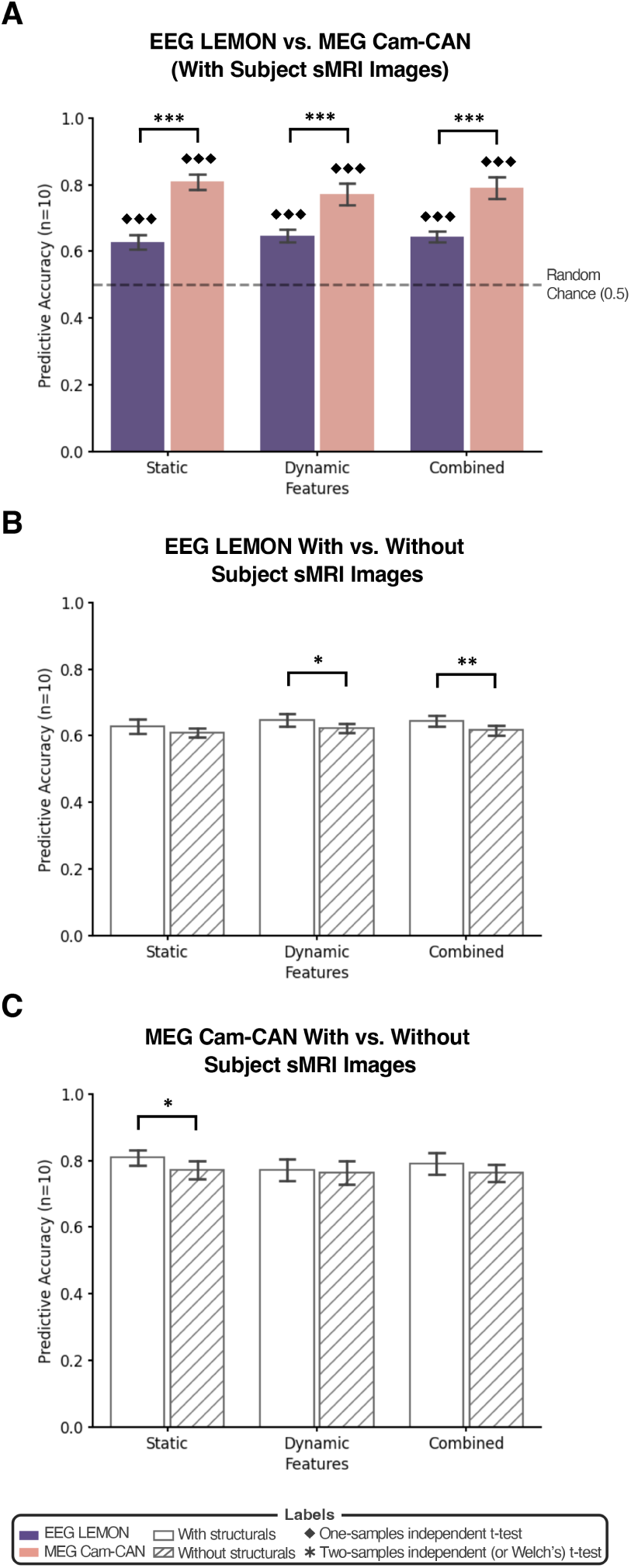
Predictive accuracy in age group classification using EEG and MEG resting-state network features. **(A)** Mean prediction scores for three different RSN feature types derived from the EEG (purple) and MEG (pink) data across 10 repeated runs of classification tasks. The bars denote mean scores, with error bars reflecting standard deviations. A dotted line represents the score expected by random chance. Features were obtained from data reconstructed using individual sMRI images. Prediction scores were evaluated against random chance with a one-sample independent t-test (for EEG, static: *T* =19.01, *p*=7.085e-9; dynamic: *T* =24.81, *p*=6.736e-10; combined: *T* =30.11, *p*=1.203e-10; for MEG, static: *T* =44.05, *p*=4.003e-12; dynamic: *T* =26.29, *p*=4.027e-10; combined: *T* =28.71, *p*=1.840e-10) and compared between EEG and MEG using a two-sample independent t-test (static: *T* =-18.47, *p*=3.801e-13; dynamic: *T* =-10.53, *p*=4.016e-9) and a Welch’s t-test (combined: *T* =-12.99, *p*=8.974e-9). **(B)** Mean prediction scores for three distinct RSN feature types derived from the EEG data that were source reconstructed with (grey) and without (orange) individual sMRI images. Bar and error bar representations match (A). Differences due to sMRI image use were assessed with a two-sample independent t-test (dynamic: *T* =3.156, *p*=5.471e-3; combined: *T* =4.076, *p*=7.088e-4). **(C)** The same analysis as in **(B)** using the MEG data (static: *T* =3.320, *p*=3.807e-3). Significance levels are indicated by */♦: *<* 0.05, **/♦♦: *<* 0.01, ***/♦♦♦: *<* 0.001 (before Bonferroni-corrected, n=3 features), with non-significant findings unmarked.

Next, we conducted the same analysis using the data that were reconstructed without subject-specific sMRI images. For EEG, the prediction scores significantly decreased when using dynamic and combined features, while a significant decrease was observed with static features for MEG (Fig. 8B, C). Irrespective of statistical significance, however, all three types of input features exhibited lower prediction scores for both EEG and MEG when only the standard MNI template was provided for reconstruction.

## 4 Discussion

Using the MEG RSNs from the Cam-CAN dataset as a benchmark for evaluating brain networks of fast oscillatory activity, we investigated EEG-derived RSNs to ascertain if both modalities can yield meaningful network descriptions and establish how comparable they are to one another. The results indicate qualitative and quantitative comparability between MEG and EEG in representing functional brain networks and suggest some advantages of HMM over traditional network inference methods for EEG (e.g., microstates [49], sliding window method [13]). Our findings highlighted several key points.

Firstly, consistent with previous research [14, 18], we found that MEG and EEG exhibit analogous static RSN features (Fig. 2), which were reproducible *within* and *across* the MEG and EEG datasets (Fig. 3). Age-related effects in static PSDs and power maps also proved comparable across the modalities. Nonetheless, there was a tendency for MEG to be more sensitive to age-related changes than EEG (Fig. 4). These network descriptions and age effects persisted even when the data were reconstructed using a standard brain structural image (Fig. A3, A4).

Secondly, MEG and EEG demonstrated broadly comparable dynamic RSN features (Fig. 5), with age-related effects in state-specific PSDs and power maps displaying marked similarity, albeit to a lesser degree compared to the static RSNs (Fig. 7). These network features and age-related effects remained largely consistent when a standard brain structural image was used to reconstruct the data (i.e., when subject-specific sMRI images were not available) (Fig. A5, A6). We also used a split-half approach to quantitatively measure the extent to which the dynamic network features were reproducible *within* and *across* the MEG and EEG datasets (Fig. 6). Overall, this approach revealed a good level of reproducibility, with state 6 (the sensorimotor network) found to be the least reproducible. We deemed a network description reproducible if the network features qualitatively resembled one another across all states, or at least the vast majority of states. We found that EEG was generally less reproducible than MEG. In sum, the similarity of dynamic RSNs across the modalities was substantial when contrasted with a previous sliding-window-based anlaysis [18] that reported discordant dynamic RSN FC between MEG and EEG.

Finally, we confirmed that both MEG and EEG can provide meaningful static and dynamic network descriptions by testing the efficacy of M/EEG network features in predicting age groups. Although EEG reported lower test accuracy than MEG, the performance scores of both modalities were above random chance expectations (Fig. 8A). However, one caveat is that, while MEG outperformed EEG in the age class prediction task, differences between the two datasets may not solely reflect differences between EEG and MEG. Other factors, such as age range and participant heterogeneity, may have also influenced the results.

Interestingly, the predictive power was similar for the static and dynamic RSN features and did not significantly improve when they were combined. This lack of improvement suggests that, for the simple task of classifying binary age groups with a logistic classifier, using either static or dynamic network descriptions suffices. The complex analysis combining static and dynamic features does not appear to be beneficial for this specific task, possibly due to the overfitting of the data. In addition, these test accuracies varied across different brain spaces. For EEG, performance metrics for dynamic and combined input features showed significant variation depending on whether the data were reconstructed with subject-specific sMRI images or the standard MNI template (Fig. 8B). For MEG, significant variation was observed in performance for static input features (Fig. 8C). Although not always statistically significant, performance metrics for all three input feature types decreased for both EEG and MEG when we used the standard MNI template.

### 4.1 Basis set of EEG RSNs

As noted at the outset of this paper, the current study provides a basis set of RSNs derived from the EEG LEMON dataset, alongside their counterparts from the MEG Cam-CAN data. This basis set was constructed using medium-density EEG, thereby rendering it as data applicable for subsequent studies employing similar EEG systems. In particular, we offer scripts designed for the preprocessing, source reconstruction, and analysis of M/EEG data, as well as the inferred model parameters of the optimal HMM runs (e.g., state covariances, transition probabilities, time course of state probabilities). These data and resources are made publicly accessible, with detailed guidelines for software installation and data downloading available in the GitHub repository (see Section 2.9).

The basis sets of static and dynamic EEG RSNs are illustrated in Figure 2 and Figure 5, respectively. The designation of these EEG networks as a basis set is justified for several reasons. Notably, in both static and dynamic analyses, EEG RSNs not only resembled those derived from the MEG data but also demonstrated age-related effects comparable across the two modalities. In addition, their network features were reproducible within and across the modalities, although with the exception of state 6 in the dynamic RSNs. Lastly, akin to MEG, network descriptions in the EEG basis set demonstrated robust performance in age prediction, significantly surpassing that of random chance.

Despite the comparability between EEG and MEG, it is important to acknowledge that MEG demonstrated better reproducibility, enhanced predictive accuracy, and greater sensitivity to age-related effects compared to EEG. This discrepancy is arguably anticipated, given the higher sensor count of MEG and consequently its superior spatial resolution. Previous studies [15, 17, 18] suggest that EEG datasets with a greater number of sensors can also achieve a level of comparability with MEG data. Nonetheless, it remains to be elucidated how EEG data, varying in sensor count relative to the EEG LEMON dataset, compares with MEG, especially in terms of dynamic RSNs as derived from state-of-the-art methodologies. Among several potential applications, this basis set could be used for investigating RSNs in high- and low-density EEG systems, developing methods for effectively combining M/EEG with fMRI, and comparing basis sets of RSNs inferred from various modalities such as optically pumped magnetometers (OPM).

### 4.2 Limitations

One of the main limitations in our study was the relatively small sample size (n=96 subjects). This limitation meant that while subjects across the two datasets could be age-matched, further matching based on sex or Mini-Mental State Examination (MMSE) scores was not feasible. Likewise, demographic attributes, recording sites, and other cognitive or physiological states that have not matched may introduced unaccounted variance into our effects of interest. In the context of our classification task, the small sample size restricted us to a binary categorisation of age groups (young vs. old), as it was difficult to subdivide our sample suitably under the k-fold cross-validation framework. Because ages were categorised in intervals within the LEMON dataset, predicting numerical, continuous age values had to be precluded from the outset. Other limitations included the use of random subsampling during the age-matching process, which may have brought additional subject variability to our comparison between LEMON and Cam-CAN.

It is also worth noting that the different data acquisition methods for EEG and MEG may have contributed to certain disparities in their RSN descriptions and associated age effects. The EEG data were recorded in 1-min blocks alternating between eyes-closed and eyes-open conditions, while the MEG data were recorded with participants keeping their eyes closed throughout the session. Although the EEG analysis was limited to the eyes-closed periods, the intermittent eye openings might have altered the brain state during EEG acquisition (e.g., by forcing participants to be more alert on average). On the contrary, the brain state in the second half of the MEG session may differ from the first half due to potential drowsiness. These differences in data acquisition likely impacted the underlying brain dynamics in EEG and MEG, leading to some disparities in the effects observed in the two modalities.

Another notable aspect of our preprocessing pipeline is that we used a parcellation atlas with 38 anatomical regions for reconstructing both EEG and MEG, which has previously been shown to work effectively on MEG data [19]. However, it is plausible that using fewer or coarser parcels could enhance the comparability between EEG and MEG. Even in this case, some differences could be preserved, for example, if localised age effects align with the adopted parcellation. Similarly, altering parcel shapes could also possibly increase comparability between the modalities if the parcel borders are defined in such a way that differences between EEG and MEG are reduced. Exploring the effects of different parcellation atlases on the EEG and MEG RSNs would be an intriguing direction for future research.

### 4.3 Outlooks and future works

Previous research has indicated marginal comparability between MEG and EEG in capturing the temporal dynamics of brain networks [18]. They proposed that the reasons for this poor correspondence are primarily two-fold. From a physiological standpoint, it was surmised that MEG and EEG detect distinct components of transient neural activities, particularly in fast oscillatory fluctuations above 1 Hz, originating from different neural populations responsible for generating magnetic and electric fields. From a methodological perspective, it was acknowledged that the sliding-window approach, compounded by the lack of spatial leakage correction, could result in large statistical errors [50]. Hence, the estimation of dynamic RSN FC at shorter timescales may not benefit from smoothing effects typically associated with static analysis techniques, such as low-pass filtering or amplitude envelope analysis.

Although we found the methodological limitations plausible, our findings did not support the physiological explanation. When we applied the TDE-HMM [20], a more advanced dynamic modelling technique validated across various MEG datasets, we discovered a previously unobserved similarity between the modalities in both static and dynamic RSNs, even with a lower-dimensional 61-channel EEG system and a 38 parcellated brain regions. Consequently, whereas it was believed that MEG and EEG produce complementary views on the dynamic functional network organisations of the human brain, our results indicate that both modalities can identify the same network profiles, at least in the resting state.

Therefore, instead of assuming they measure distinct aspects of brain functionality [51, 52], our data suggest that the same functional networks underpin brain activity, irrespective of the modalities. This indicates that the RSNs may be less a characteristic of the specific measurement modality and more a neural substrate correlated with cognition and behaviours that is common across neuroimaging modalities. It is conceivable that different modalities may highlight specific aspects of RSNs, as evidenced by statistical testing on age effects; however, the overarching spatial patterns remained consistent. With the application of robust statistical methods capable of enhancing spatiotemporal precision [53], such as Dynamic Network Modes (DyNeMo) [5], we may be able to see more clearly the same RSNs in both MEG and EEG.

Our results thus support the notion that previously known properties of MEG and fMRI RSNs may translate into EEG. Furthermore, while both MEG and EEG provide unprecedented temporal resolution over fMRI, their comparability allow us to consider prioritising EEG in future functional brain network studies, given its cheap cost and broader accessibility. If high-density EEG systems are leveraged in conjunction with advanced statistical modelling methodologies, we anticipate an even greater alignment in the dynamic network features identifiable across these modalities.

To this extent, future works should aim to verify whether cognitive, behavioural, and clinical phenotypes associated with MEG and fMRI RSNs can also be identified in EEG. Given that dynamic RSN phenotypes of neuropsychiatric disorders have predominantly been reported in fMRI and MEG studies, it would be crucial to explore dynamic EEG RSNs in this regard. The established correspondence between MEG and fMRI RSNs [3] implies that biomarkers identified through these modalities could potentially be detected within EEG data. An additional avenue of research could focus on the use of simultaneous M/EEG recordings to more accurately assess how dynamic RSN features and the age-related effects in them vary between modalities. This approach would offer insights into the efficacy and necessity of multimodal electrophysiological techniques. Furthermore, examining whether the comparability of RSNs across MEG, EEG, and fMRI extends to other measurement devices, such as wearable functional near-infrared spectroscopy (fNIRS) [54] or OPM-based MEG systems [55], would substantiate the validity of our findings.

## 5 Conclusion

In this paper, we demonstrated that MEG and EEG produce comparable static and dynamic network descriptions, which were shown to be meaningful in highlighting age-related effects and predicting age groups. MEG, however, was more reproducible and sensitive in detecting age-related effects than EEG. This comparability and concomitant discrepancies between the modalities were also observed when the datasets were reconstructed using a standard brain structural image and without requiring subject sMRI images, albeit to a lesser degree. Our results suggest that MEG and EEG share the same functional brain networks, likely paralleling those detected by other modalities like fMRI.

## 7 Credit Authorship Contribution Statement

SC: Conceptualisation, Data curation, Software, Methodology, Formal analysis, Writing – original draft, Writing – review and editing; ME: Conceptualisation, Software, Writing – review and editing, Supervision; MW: Conceptualisation, Methodology, Writing – review and editing, Supervision; CG: Conceptualisation, Data curation, Software, Methodology, Writing – review and editing, Supervision.

## Acknowledgements

MW is supported by the Wellcome Trust (106183/Z/14/Z, 215573/Z/19/Z), the New Therapeutics in Alzheimer’s Diseases (NTAD) study supported by UK MRC, the Dementia Platform UK (RG94383/RG89702), and the NIHR Oxford Health Biomedical Research Centre (NIHR203316). The views expressed are those of the author(s) and not necessarily those of the NIHR or the Department of Health and Social Care.

## Appendix A Supplementary Figures

**Fig. A1.**
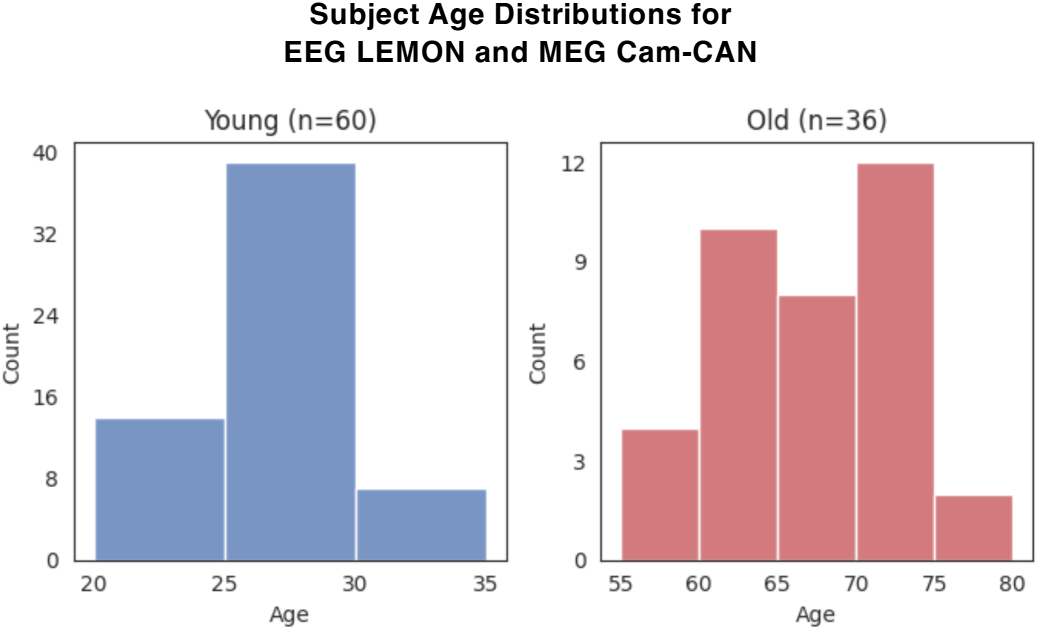
Age distributions of participants in the EEG LEMON and MEG Cam-CAN datasets. Histograms depict the distributions of ages for the young (blue) and old (red) group. Note that the EEG and MEG data have identical distributional shapes, as the subjects in two datasets are age-matched. To ensure comparable sample sizes between LEMON and Cam-CAN, a subset of participants was sampled from each age group for each dataset. Both datasets have a sample size of 60 young and 36 old participants.

**Fig. A2.**
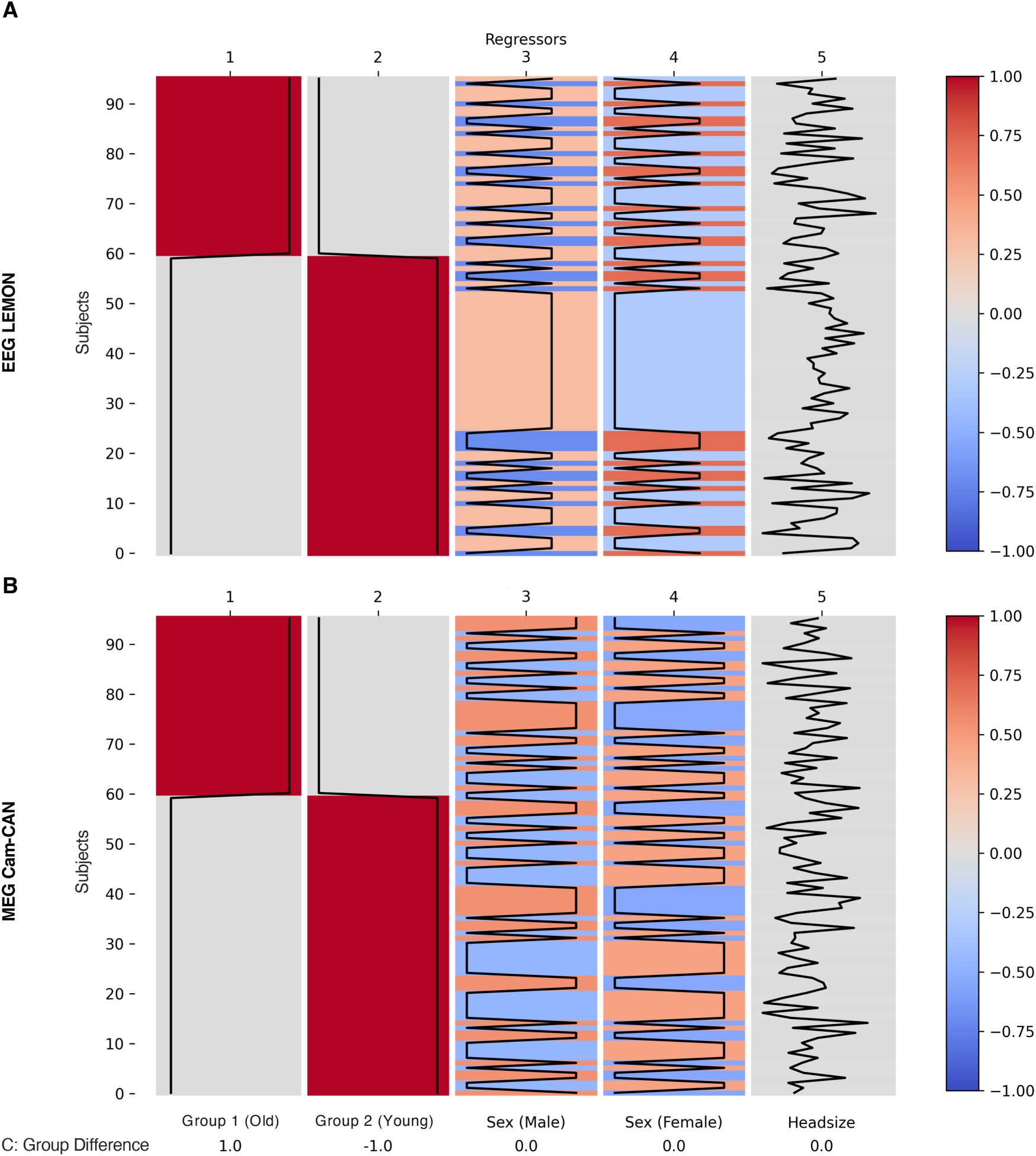
Group-level design matrices and contrasts for the EEG and MEG datasets. **(A)** The design matrix for the EEG data with regressors in columns and individual subjects in rows. The matrix consists of the condition (Group) and covariate (Sex, Headsize) regressors, with covariate values demeaned. A colour bar indicates the values for each regressor. The table below outlines the contrast (i.e., mean group difference), denoted as C, including the weightings associated with each regressor. **(B)** The design matrix for the MEG data visualised in the same structure as in (A).

**Fig. A3.**
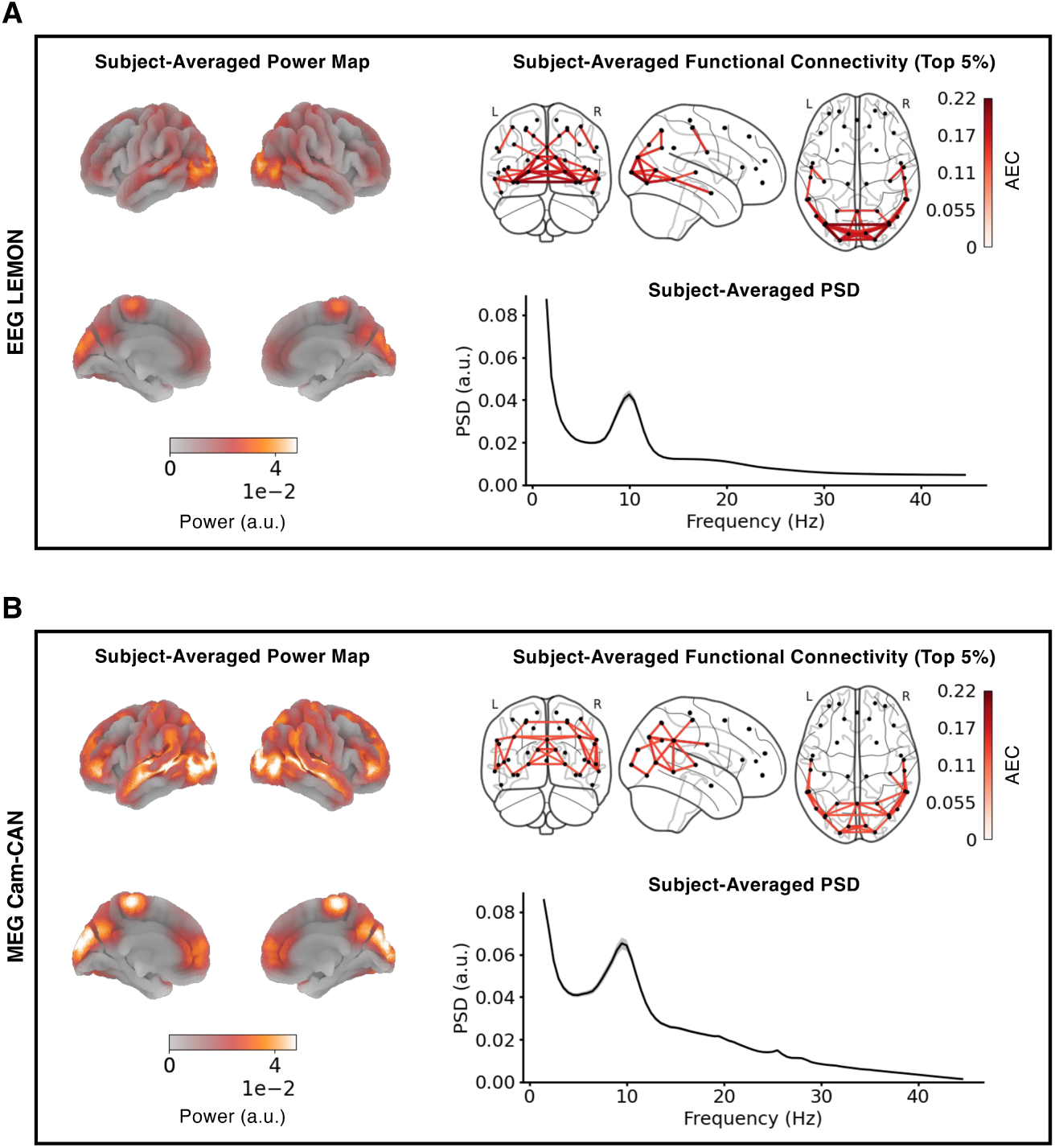
Static resting-state networks inferred from datasets source reconstructed using a standard brain structural image. **(A)** A subject-averaged, wide-band (1.5-20 Hz) power map displayed for the EEG LEMON dataset (left). On the top right, a subject-averaged, wide-band (1.5-20 Hz) FC network, with only the top 5% of correlation strengths shown for visualisation purposes. On the bottom right, a parcel- and subject-averaged PSD, with standard errors across parcels indicated by grey shading. **(B)** The subject-averaged power map, FC network, and PSD are generated using the MEG Cam-CAN dataset following the same analyses described in (A). For both EEG and MEG, the data were source reconstructed using a standard MNI brain, not individual sMRI files.

**Fig. A4.**
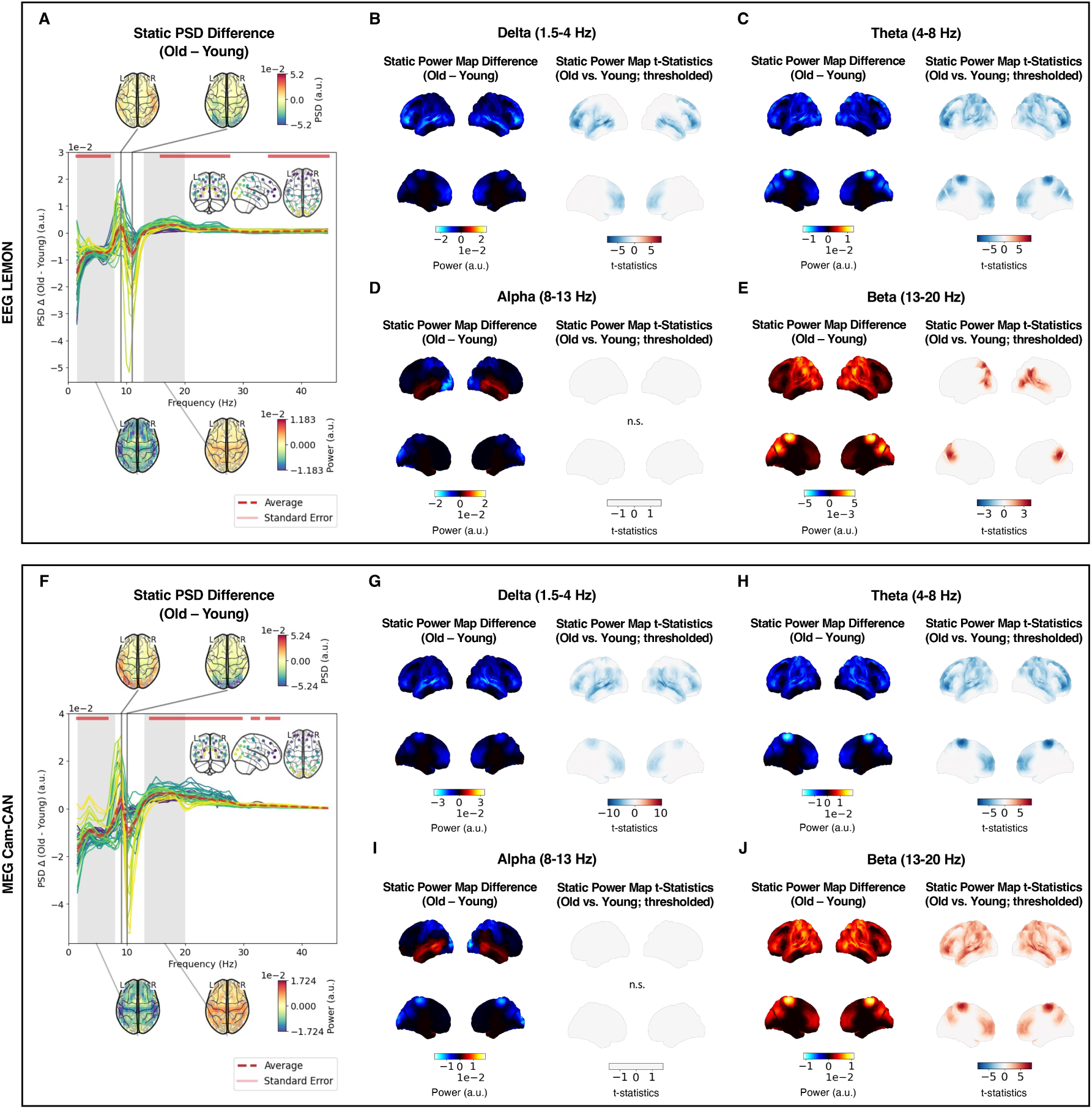
Age-related effects in static network features extracted using a standard brain structural image. **(A)** For the EEG data, the between-group PSD difference (red dotted line), averaged over parcels and subjects, and its standard error (pink shade) are depicted alongside green lines denoting the parcel-wise, subject-averaged PSD differences. An inset in the top-right corner marks the brain parcel locations associated with each PSD difference line. Frequency ranges with significant (p *<* 0.05) PSD differences between the two age groups are indicated by red bars. Additionally, the group-level peak differences in the alpha (8-13 Hz) band and group-level PSD differences integrated across the lower (1.5-8 Hz) and beta (13-20 Hz) bands are depicted in topographical maps at the top and bottom, respectively. **(B)** On the left, a brain surface map illustrating delta band power differences between the old and young groups is plotted for the EEG data. On the right, the brain regions with significant (p *<* 0.05; Bonferroni-corrected, n=4 frequency bands) between-group differences are highlighted. Red and blue colours indicate higher power in old and young participants, respectively, and non-significant findings are labelled as n.s. The same analysis as in (B) was replicated for the **(C)** theta, **(D)** alpha, and **(E)** beta band EEG power maps. **(F)** The same analysis as described in (A) is conducted on the MEG data. **(G)-(J)** The same analysis as described in (B) is applied to delta, theta, alpha, and beta band power differences using the MEG data, respectively. For both EEG and MEG, the data were source reconstructed using a standard MNI structural brain image.

**Fig. A5.**
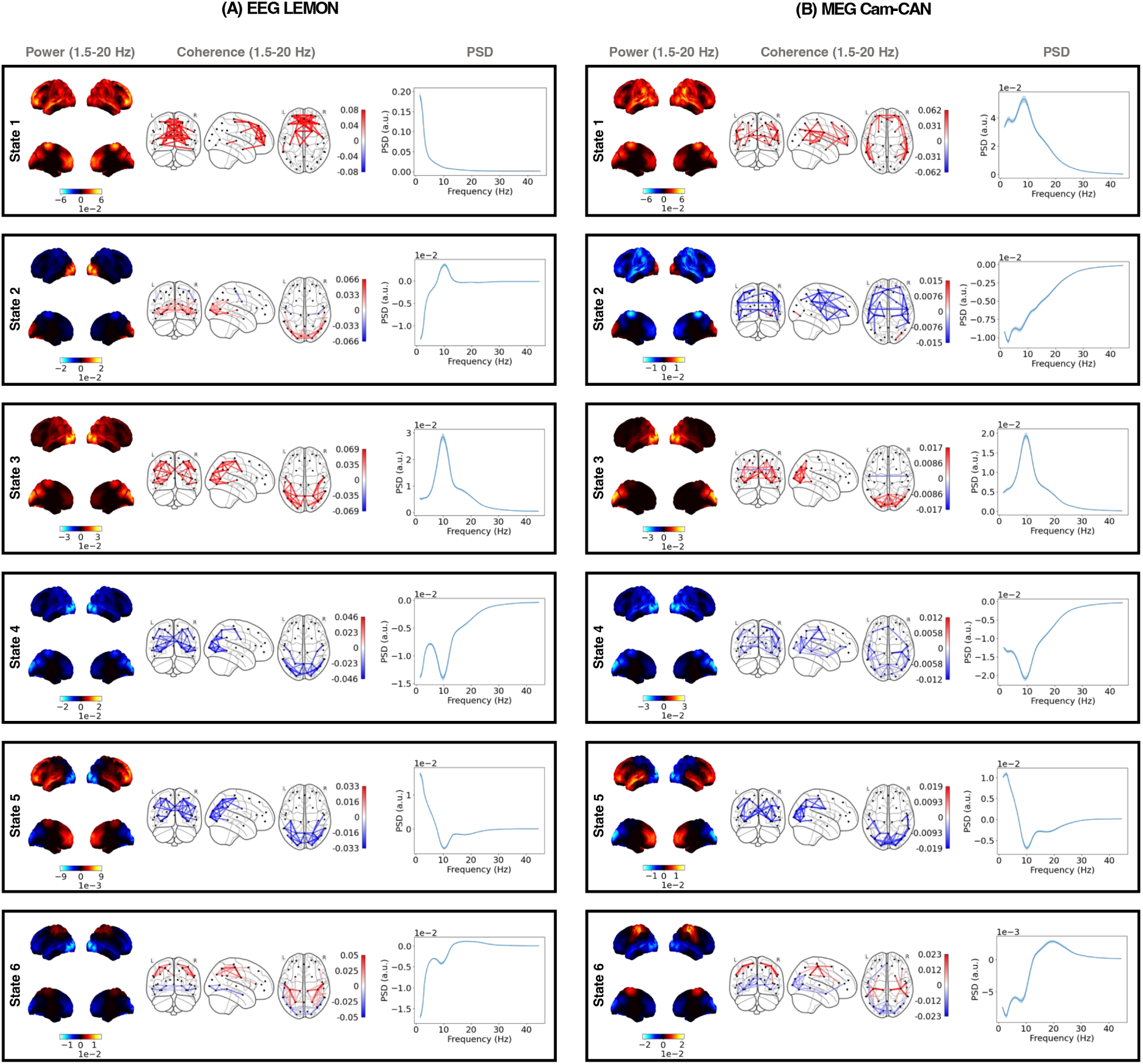
Dynamic resting-state networks inferred from datasets source reconstructed using a standard brain structural image. **(A)** Each box shows the subject-averaged power map (left), FC network (middle), and parcel-averaged PSD (right) of each HMM state for 96 EEG subjects. The power maps show lateral surfaces at the top and medial surfaces at the bottom. The FC networks illustrate connections with the top 5% coherence values. The shaded areas of the PSDs represent the standard error of the mean. The power maps, FC networks, and PSDs are visualised relative to their average across all states. **(B)** The plots follow the same format as (A), showing the RSNs computed from 96 MEG subjects. For both EEG and MEG, the data were source reconstructed using a standard MNI brain and not individual sMRI files.

**Fig. A6.**
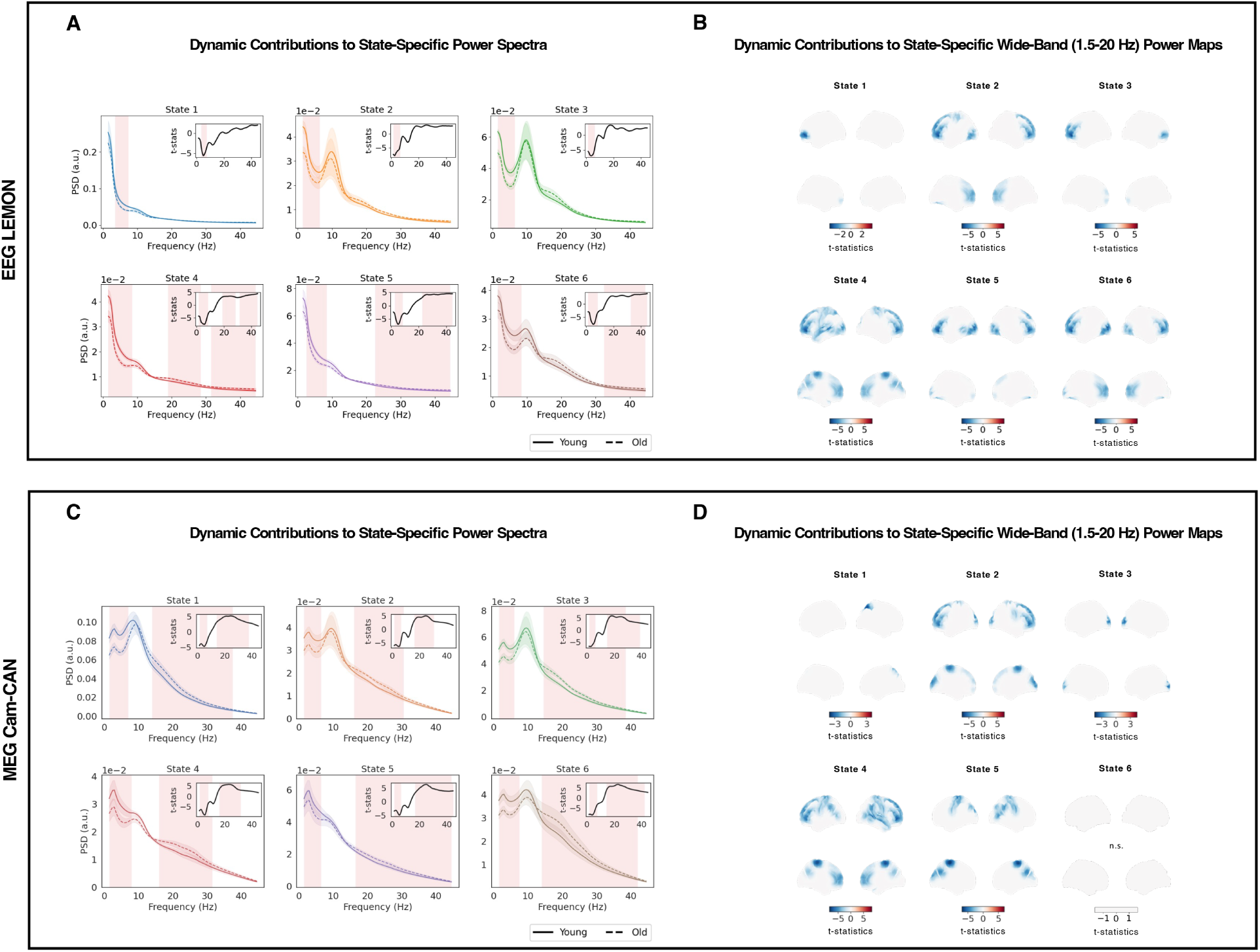
Age-related effects in dynamic network features extracted using a standard brain structural image. **(A)** The state-specific PSDs of the EEG data are averaged across grouped subjects and parcels and depicted for old (dotted) and young (solid) participants. For each HMM state, frequencies exhibiting statistically significant (p *<* 0.05; Bonferroni-corrected, n=6 states) age-related dynamic effects are highlighted in red. An inset at the top-right corner displays the t-statistics, signifying the magnitude of age-related effect sizes between the groups. **(B)** The state-specific power maps with significant (p *<* 0.05; Bonferroni-corrected, n=6 states) between-group differences are shown for the EEG data. Red and blue colours indicate higher power in old and young participants, respectively, and non-significant findings are labelled as n.s. **(C)** The same analysis as in (A) is performed on the MEG data, marking significant age-related dynamic effects in state-specific PSDs. **(D)** The same analysis as in (B) is performed on the MEG data, illustrating significant age-related dynamic effects in state-specific power maps. For both EEG and MEG, the data were source reconstructed using a standard MNI structural brain image.

**Fig. A7.**
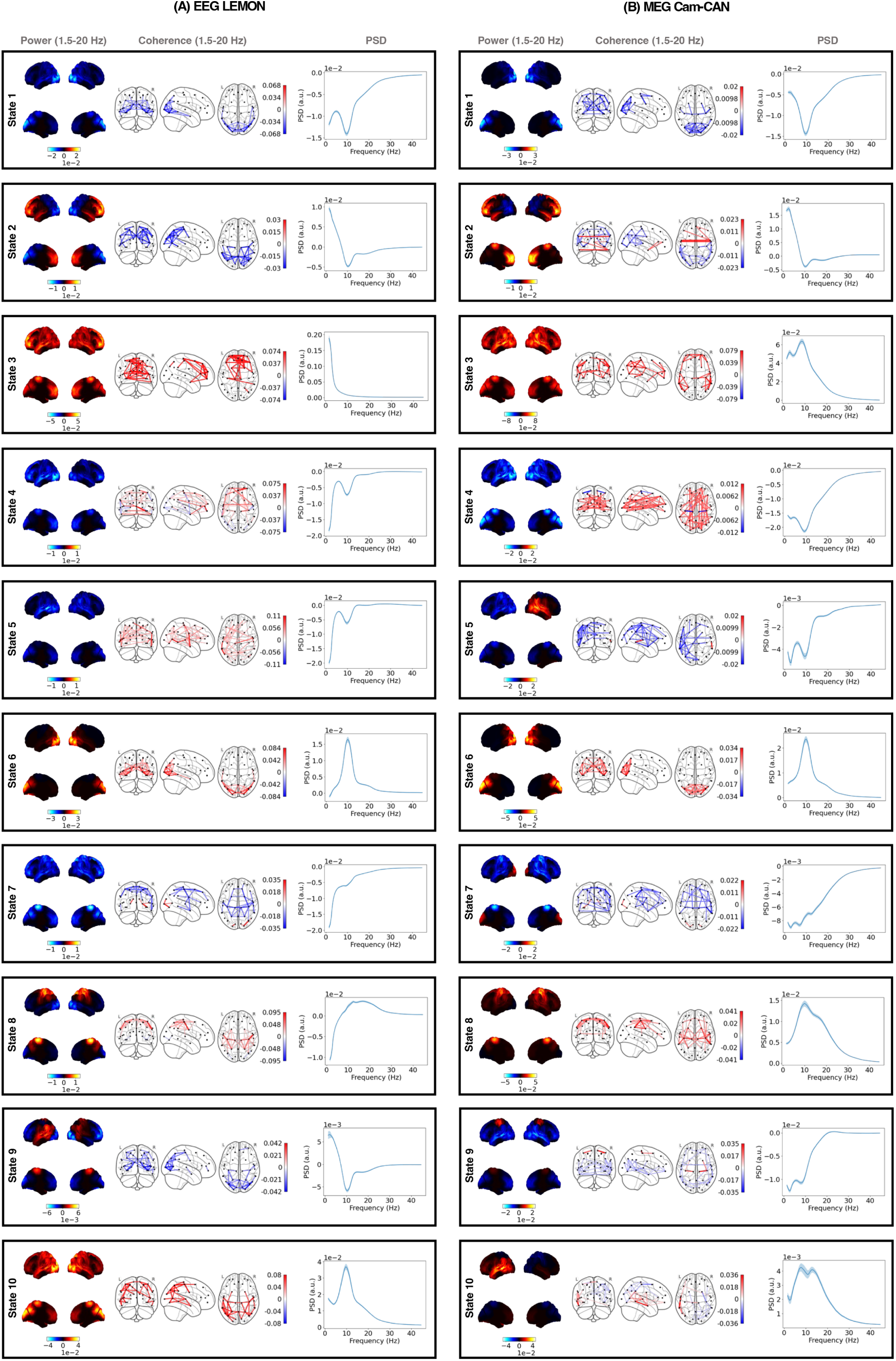
10-state dynamic resting-state networks of EEG and MEG. Dynamic RSNs of 96 subjects are shown for the (A) EEG and (B) MEG datasets, identical to Figure 5, but with 10 HMM states inferred from the data. Note that source reconstruction was performed using individual sMRI images for both EEG and MEG.

## Footnotes

1 These age ranges were selected for the young and old cohorts based on the age distribution of the LEMON dataset (i.e., how the two groups were divided in the original dataset) [21].

2 It is worth considering that, in matching the rank of both the MEG and EEG data to 50, we may have lost some of the additional spatial resolution available in MEG, as the MEG data has a higher rank than EEG data at the sensor level, even after maxfiltering. Therefore, this matched rank could have artificially increased the similarity between the EEG and MEG descriptions beyond what would naturally occur. In this paper, however, we decided that the most appropriate strategy is to align the preprocessing steps as closely as possible across both modalities, to ensure that any observed differences between the two are not due to trivial variations arising from preprocessing and source reconstruction steps.

3 This procedure applied exclusively to analyses involving data reconstructed with subject sMRI images. For data reconstructed with the standard MNI template, the procedure was not reiterated. Instead, the optimal run was chosen from a total of 10 runs.

4 Typically, state alignments in multiple HMM runs are conducted automatically through the linear sum assignment algorithm [36]. This approach tends to yield satisfactory results only when HMMs are trained on datasets of identical modalities. Aligning states across HMM runs trained on datasets of different modalities thus necessitated manual intervention due to the algorithm’s decreased efficacy under such conditions. However, it should be noted that while manual alignment by eye was reliable for matching six states in our case, this process may still have introduced some human-induced bias.

## References

[1] Biswal B, Zerrin Yetkin F, Haughton VM, Hyde JS. Functional connectivity in the motor cortex of resting human brain using echo-planar mri. Magnetic Resonance in Medicine. 1995;34(4):537–541. 10.1002/mrm.1910340409.

[2] Shulman GL, Fiez JA, Corbetta M, Buckner RL, Miezin FM, Raichle ME, et al. Common Blood Flow Changes across Visual Tasks: II. Decreases in Cerebral Cortex. Journal of Cognitive Neuroscience. 1997;9(5):648–663. 10.1162/jocn.1997.9.5.648.

[3] Brookes MJ, Woolrich M, Luckhoo Hy, Price D, Hale JR, Stephenson MC, et al. Investigating the electrophysiological basis of resting state networks using magnetoencephalography. Proceedings of the National Academy of Sciences. 2011;108(40):16783–16788. 10.1073/pnas.1112685108.

[4] Baker AP, Brookes MJ, Rezek IA, Smith SM, Behrens T, Probert Smith PJ, et al. Fast transient networks in spontaneous human brain activity. eLife. 2014;3:e01867. 10.7554/elife.01867.

[5] Gohil C, Roberts E, Timms R, Skates A, Higgins C, Quinn A, et al. Mixtures of large-scale dynamic functional brain network modes. NeuroImage. 2022;263:119595. 10.1016/j.neuroimage.2022.119595.

[6] Quinn AJ, Vidaurre D, Abeysuriya R, Becker R, Nobre AC, Woolrich MW. Task-Evoked Dynamic Network Analysis Through Hidden Markov Modeling. Frontiers in Neuroscience. 2018;12:603. 10.3389/fnins.2018.00603.

[7] Barnes JJ, Woolrich MW, Baker K, Colclough GL, Astle DE. Electrophysiological measures of resting state functional connectivity and their relationship with working memory capacity in childhood. Developmental Science. 2015;19(1):19–31. 10.1111/desc.12297.

[8] Becker R, Vidaurre D, Quinn AJ, Abeysuriya RG, Parker Jones O, Jbabdi S, et al. Transient spectral events in resting state MEG predict individual task responses. NeuroImage. 2020;215:116818. 10.1016/j.neuroimage.2020.116818.

[9] Sitnikova TA, Hughes JW, Ahlfors SP, Woolrich MW, Salat DH. Short timescale abnormalities in the states of spontaneous synchrony in the functional neural networks in Alzheimer’s disease. NeuroImage: Clinical. 2018;20:128–152. 10.1016/j.nicl.2018.05.028.

[10] Leuchter AF, Cook IA, Hunter AM, Cai C, Horvath S. Resting-State Quantitative Electroencephalography Reveals Increased Neurophysiologic Connectivity in Depression. PLoS ONE. 2012;7(2):e32508. 10.1371/journal.pone.0032508.

[11] Lottman KK, Gawne TJ, Kraguljac NV, Killen JF, Reid MA, Lahti AC. Examining resting-state functional connectivity in first-episode schizophrenia with 7T fMRI and MEG. NeuroImage: Clinical. 2019;24:101959. 10.1016/j.nicl.2019.101959.

[12] Mantini D, Perrucci MG, Del Gratta C, Romani GL, Corbetta M. Electrophysiological signatures of resting state networks in the human brain. Proceedings of the National Academy of Sciences. 2007;104(32):13170–13175. 10.1073/pnas.0700668104.

[13] de Pasquale F, Della Penna S, Snyder AZ, Lewis C, Mantini D, Marzetti L, et al. Temporal dynamics of spontaneous MEG activity in brain networks. Proceedings of the National Academy of Sciences. 2010;107(13):6040–6045. 10.1073/pnas.0913863107.

[14] Siems M, Pape A, Hipp JF, Siegel M. Measuring the cortical correlation structure of spontaneous oscillatory activity with EEG and MEG. NeuroImage. 2016;129:345–355. 10.1016/j.neuroimage.2016.01.055.

[15] Knyazev GG, Savostyanov AN, Bocharov AV, Tamozhnikov SS, Saprigyn AE. Task-positive and task-negative networks and their relation to depression: EEG beamformer analysis. Behavioural Brain Research. 2016;306:160–169. 10.1016/j.bbr.2016.03.033.

[16] Sockeel S, Schwartz D, Peĺegrini-Issac M, Benali H. Large-Scale Functional Networks Identified from Resting-State EEG Using Spatial ICA. PLoS ONE. 2016;11(1):e0146845. 10.1371/journal.pone.0146845.

[17] Liu Q, Farahibozorg S, Porcaro C, Wenderoth N, Mantini D. Detecting large-scale networks in the human brain using high-density electroencephalography: Imaging Brain Networks with High Density EEG. Human Brain Mapping. 2017;38(9):4631–4643. 10.1002/hbm.23688.

[18] Coquelet N, De Tìege X, Destoky F, Roshchupkina L, Bourguignon M, Goldman S, et al. Comparing MEG and high-density EEG for intrinsic functional connectivity mapping. NeuroImage. 2020;210:116556. 10.1016/j.neuroimage.2020.116556.

[19] Colclough G, Brookes M, Smith S, Woolrich M. A symmetric multivariate leakage correction for MEG connectomes. NeuroImage. 2015;117:439–448. 10.1016/j.neuroimage.2015.03.071.

[20] Vidaurre D, Hunt LT, Quinn AJ, Hunt BAE, Brookes MJ, Nobre AC, et al. Spontaneous cortical activity transiently organises into frequency specific phase- coupling networks. Nature Communications. 2018;9(1):2987. 10.1038/s41467-018-05316-z.

[21] Babayan A, Erbey M, Kumral D, Reinelt JD, Reiter AMF, Robbig J, et al. A mind-brain-body dataset of MRI, EEG, cognition, emotion, and peripheral physiology in young and old adults. Scientific Data. 2019;6(1):180308. 10.1038/sdata.2018.308.

[22] Shafto MA, Tyler LK, Dixon M, Taylor JR, Rowe JB, Cusack R, et al. The Cambridge Centre for Ageing and Neuroscience (Cam-CAN) study protocol: a cross-sectional, lifespan, multidisciplinary examination of healthy cognitive ageing. BMC Neurology. 2014;14:204. 10.1186/s12883-014-0204-1.

[23] Gohil C, Huang R, Roberts E, van Es MWJ, Quinn AJ, Vidaurre D, et al. osl-dynamics: A toolbox for modelling fast dynamic brain activity. eLife. 2023;12:RP91949. 10.7554/eLife.91949.2.

[24] Quinn AJ, van Es MW, Gohil C, Woolrich MW. OHBA Software Library in Python (OSL) (0.1.1). Zenodo. 2023;10.5281/zenodo.6875059.

[25] Rosner B. Percentage Points for a Generalized ESD Many-Outlier Procedure. Technometrics. 1983;25:165–172. 10.2307/1268549.

[26] Hyvarinen A. Fast and robust fixed-point algorithms for independent component analysis. IEEE Transactions on Neural Networks. 1999;10:626–634. 10.1109/72.761722.

[27] Perrin F, Pernier J, Bertrand O, Echallier J. Spherical splines for scalp potential and current density mapping. Electroencephalography and Clinical Neurophysiology. 1989;72:184–187. 10.1016/0013-4694(89)90180-6.

[28] Smith SM. Fast robust automated brain extraction. Human Brain Mapping. 2002;17(3):143–155. 10.1002/hbm.10062.

[29] Jenkinson M, Pechaud M, Smith S.: BET2: MR-based estimation of brain, skull and scalp surfaces. In Eleventh annual meeting of the organization for human brain mapping, 17:167. Toronto, Canada.

[30] Veen BV, Buckley K. Beamforming: a versatile approach to spatial filtering. IEEE ASSP Magazine. 1988;5(2):4–24. 10.1109/53.665.

[31] Palva JM, Wang SH, Palva S, Zhigalov A, Monto S, Brookes MJ, et al. Ghost interactions in MEG/EEG source space: A note of caution on inter-areal coupling measures. NeuroImage. 2018;173:632–643. 10.1016/j.neuroimage.2018.02.032.

[32] Bishop CM. Pattern Recognition and Machine Learning. New York: Springer; 2006.

[33] Rezek I, Roberts S. Ensemble Hidden Markov Models with Extended Observation Densities for Biosignal Analysis. In: Husmeier D, Dybowski R, Roberts S, editors. Probabilistic Modeling in Bioinformatics and Medical Informatics. London: Springer; 2005. p. 419–450.

[34] Vidaurre D, Abeysuriya R, Becker R, Quinn AJ, Alfaro-Almagro F, Smith SM, et al. Discovering dynamic brain networks from big data in rest and task. NeuroImage. 2018;180:646–656. 10.1016/j.neuroimage.2017.06.077.

[35] Kingma DP, Ba J. Adam: A method for stochastic optimization. arXiv. 2014;p. 1412.6980. 10.48550/arXiv.1412.6980.

[36] Crouse DF. On implementing 2D rectangular assignment algorithms. IEEE Transactions on Aerospace and Electronic Systems. 2016;52(4):1679–1696. 10.1109/TAES.2016.140952.

[37] Quinn AJ, Atkinson LZ, Gohil C, Kohl O, Pitt J, Zich C, et al. The GLM-spectrum: A multilevel framework for spectrum analysis with covariate and confound modelling. Imaging Neuroscience. 2024;2:1–26. 10.1162/imaga00082.

[38] Welch P. The use of fast Fourier transform for the estimation of power spectra: A method based on time averaging over short, modified periodograms. IEEE Transactions on Audio and Electroacoustics. 1967;15(2):70–73. 10.1109/TAU.1967.1161901.

[39] Vidaurre D, Quinn AJ, Baker AP, Dupret D, Tejero-Cantero A, Woolrich MW. Spectrally resolved fast transient brain states in electrophysiological data. NeuroImage. 2016;126:81–95. 10.1016/j.neuroimage.2015.11.047.

[40] Colclough GL, Woolrich MW, Tewarie PK, Brookes MJ, Quinn AJ, Smith SM. How reliable are MEG resting-state connectivity metrics? NeuroImage. 2016;138:284–293. 10.1016/j.neuroimage.2016.05.070.

[41] Vidaurre D, Llera A, Smith SM, Woolrich MW. Behavioural relevance of spontaneous, transient brain network interactions in fMRI. NeuroImage. 2021;229:117713. 10.1016/j.neuroimage.2020.117713.

[42] Huang R, Gohil C, Woolrich M. Modelling variability in dynamic functional brain networks using embeddings. bioRxiv. 2024;10.1101/2024.01.29.577718.

[43] Holmes AP, Blair RC, Watson JDG, Ford I. Nonparametric Analysis of Statistic Images from Functional Mapping Experiments. Journal of Cerebral Blood Flow & Metabolism. 1996;16(1):7–22. 10.1097/00004647-199601000-00002.

[44] Nichols TE, Holmes AP. Nonparametric permutation tests for functional neuroimaging: A primer with examples. Human Brain Mapping. 2002;15:1–25. 10.1002/hbm.1058.

[45] Robert P, Escoufier Y. A Unifying Tool for Linear Multivariate Statistical Methods: The RV-Coefficient. Applied Statistics. 1976;25(3):257. 10.2307/2347233.

[46] Lin J. Divergence measures based on the Shannon entropy. IEEE Transactions on Information Theory. 1991;37(1):145–151. 10.1109/18.61115.

[47] Ng AY.: Feature selection, L1 vs. L2 regularization, and rotational invariance. In Proceedings of the 21st International Conference on Machine Learning. Banff, Canada.

[48] Hoerl AE, Kennard RW. Ridge Regression: Biased Estimation for Nonorthogonal Problems. Technometrics. 1970;12(1):55. 10.2307/1267351.

[49] Michel CM, Koenig T. EEG microstates as a tool for studying the temporal dynamics of whole-brain neuronal networks: A review. NeuroImage. 2018;180:577–593. 10.1016/j.neuroimage.2017.11.062.

[50] Hutchison RM, Womelsdorf T, Allen EA, Bandettini PA, Calhoun VD, Corbetta M, et al. Dynamic functional connectivity: Promise, issues, and interpretations. NeuroImage. 2013;80:360–378. 10.1016/j.neuroimage.2013.05.079.

[51] Cichy RM, Pantazis D. Multivariate pattern analysis of MEG and EEG: A comparison of representational structure in time and space. NeuroImage. 2017;158:441–454. 10.1016/j.neuroimage.2017.07.023.

[52] Mahmutoglu MA, Rupp A, Baumgartner U. Simultaneous EEG/MEG yields complementary information of nociceptive evoked responses. Clinical Neurophysiology. 2022;143:21–35. 10.1016/j.clinph.2022.08.005.

[53] McFadyen J, Dolan RJ. Spatiotemporal Precision of Neuroimaging in Psychiatry. Biological Psychiatry. 2023;93:671–680. 10.1016/j.biopsych.2022.08.016.

[54] Ferrari M, Quaresima V. A brief review on the history of human functional near-infrared spectroscopy (fNIRS) development and fields of application. NeuroImage. 2012;63(2):921–935. 10.1016/j.neuroimage.2012.03.049.

[55] Brookes MJ, Leggett J, Rea M, Hill RM, Holmes N, Boto E, et al. Magnetoencephalography with optically pumped magnetometers (OPM-MEG): the next generation of functional neuroimaging. Trends in Neuroscience. 2022;45(8):621–634. 10.1016/j.tins.2022.05.008.

